# Color-specific porosity in double pigmented natural 3d-nanoarchitectures of blue crab shell

**DOI:** 10.1101/762948

**Authors:** Fran Nekvapil, Simona Cintă Pinzaru, Lucian Barbu–Tudoran, Maria Suciu, Branko Glamuzina, Tudor Tamaş, Vasile Chiş

## Abstract

3D-engineered nanoarchitectures with various functionalities are still difficult to obtain and translate for real-world applications. However, such nanomaterials are naturally abundant and yet wasted, but could trigger huge interest for blue bioeconomy, provided that our understanding of their ultrastructure-function is achieved. To date, the Bouligand pattern in crustaceans shell structure is believed to be unique. Here we demonstrated that in blue crab *Callinectes sapidus*, the 3D-nanoarchitecture is color-specific, while the blue and red-orange pigments interplay in different nano-sized channels and pores. Thinnest pores of about 20 nm are found in blue shell. Additionally, the blue pigment co-existence in specific Bouligand structure is proved for the green crab *Carcinus aestuarii*, although the crab does not appear blue. The pigments interplay, simultaneously detected in color-specific, native crab shells overturns our understanding in crustaceans coloration and may trigger the selective use of particular colored natural nanoarchitectures for biomolecular loading and slow release, infectious barriers, bioremediation, translational diffusivity and many others.

---

Nikola Tesla would have been surprised to find out that thinking “in terms of energy, frequency, and vibration”, the secrets of the blue crab colors could be elucidated.

Accounted by ecologists among 100 worst invasive alien species in eastern Mediterranean coast^1–5^, seen by seafood producers both as a threat to benthic shellfish cultures^2,4–6^ and a potential new commodity in the invaded areas^4,6^, loved by gourmands as a delicacy, the Atlantic blue crab, *Callinectes sapidus*, poses an increasing interest, as its wasted shells could be potentially turned in valuable by-product, not only because of chitin, proteins^7^ and biogenic calcite content, but also as natural, porous biomaterial, yet poorly understood as highly ordered nanoarchitecture posing inspiration for biomimetics^8^.

Interestingly, *C. sapidus* cuticle simultaneously features bright blue, red and white anatomical exoskeleton counterparts (Supplementary Fig. 1), known to be important cues for mate selection for reproduction^9^. However, considering the large diversity of crustaceans’ coloration, research on blue color origin refers mostly to lobster species *Homarus americanus*^10–13^ and *Homarus gamarus*^14–19^ which is presumed to originate from blue carotenoprotein complexes, where astaxanthin (ATX), an orange carotenoid is non-covalently bonded in crustacyanins^10–21^. UV/Vis absorption band shifted from 480 nm in ATX monomers^12^ to 632, 585 - 595 and 610 nm in isolated α-, β- and γ-crustacyanin^12^ respectively, suggests the opportunity to exploit vibrational resonance Raman (RR) spectroscopy to selectively detect free or non-covalently bound ATX in custacyanin in intact crab shell by-product.

It is also surprising that the current knowledge on the blue coloration in crustaceans remains controversial, considering the numerous earlier studies^10,11,12,14,15^, and more recent computational and experimental approaches^13,16–19^, ^22–26^. For instance, Gamiz-Hernandez et al^19^ showed that β-crustacyanin (not α-) is the responsible pigment for the blue color. Based on theoretical calculations^19^, it was found that β-crustacyanin (comprising two stacked ATX molecules), induced a bathochromic shift of ATX arising from the polarization effects and steric constrains of the ATX-protein binding site. On the other hand, based on the available models of crustacyanins, van Wijk et al.^17^ proposed a protonation model involving the keto-groups of the ATX terminals near a water molecule. Controversial conclusions underlying the comparative results of the cuticle ultrastructure in lobster *H. americanus* raised additional questions regarding its crystalline nature^27,28^. The “universal” nature of the 3D- hierarchical calcite arrays which follows a twisting plywood path (Bouligand-type pattern) in crustacean exoskeleton has been suggested, based on studies on several species of crabs^8,28–32^, but lacking any connection with the pigments populating the morphological pattern. Katsikini^33^ detected ATX only in *C. sapidus* blue crab cuticle, due to the use of a single laser line (488 nm), which fulfilled the RR conditions for the respective carotenoid only, but suggested that Br and Sr are involved in two different mechanisms contributing to the blue color. Currently, comprehensive understanding of chemical structure-morphology relationship in blue crab shell is absent, although of high interest for transforming such aquatic porous by-product into added-value material within the blue bioeconomy goal.

Here, relying on combinatory multi-laser RR micro-spectroscopy and imaging, assisted by electron microscopy, X-ray diffraction and computational chemistry, we show that two interplayed pigments and their distribution in the 3D-nanoarchitecture of the plywood path is color-specific and the blue shell of *C. sapidus* is not exclusively determined by the presence blue pigment, astaxanthin bounded in crustacyanin. Moreover the blue pigment is present in other crab cuticle colors, not necessarily blue. Comparatively, we employed a co-inhabitant, native^2^ species, *Carcinus aestuarii* green crab. The ultimate goal is to properly address such wasted biogenic material within the blue bioeconomy concept.

We hypothesize an additional photonic color component to the overall blue cuticle appearance, sustained by its optical properties^9,33^ (high reflectance between 380-480 nm, selective absorption^33^ between 500-700 nm, weak light transmission of about 9.6% of red *C. sapidus* shell^33^) and, yet limited, RR scattering data of one blue shell^33^. Exoskeleton photonic crystal would be sustained, provided that the ultrastructure periodicity influences the propagation of certain light wavelengths and its dielectric properties are tuned by its morphological biochemistry.

## Results

The complete experimental approach to extract information on pigments identity and their native distribution in crabs exoskeleton ultrastructure is summarized in the Fig. 1. We correlate the unique ability of the multi-laser (resonance) Raman spectroscopic techniques to localize at sub-micrometer level the structural information with the morphology from the high resolution scanning electron microscopy (HR-SEM) in conjunction with energy-dispersive X-ray spectroscopy (EDX) and X-ray powder diffraction (XRD) in native blue, red and white counterparts of *C. sapidus* claw shells, along with the green *C. aestuarii* crab.

**Fig. 1.**
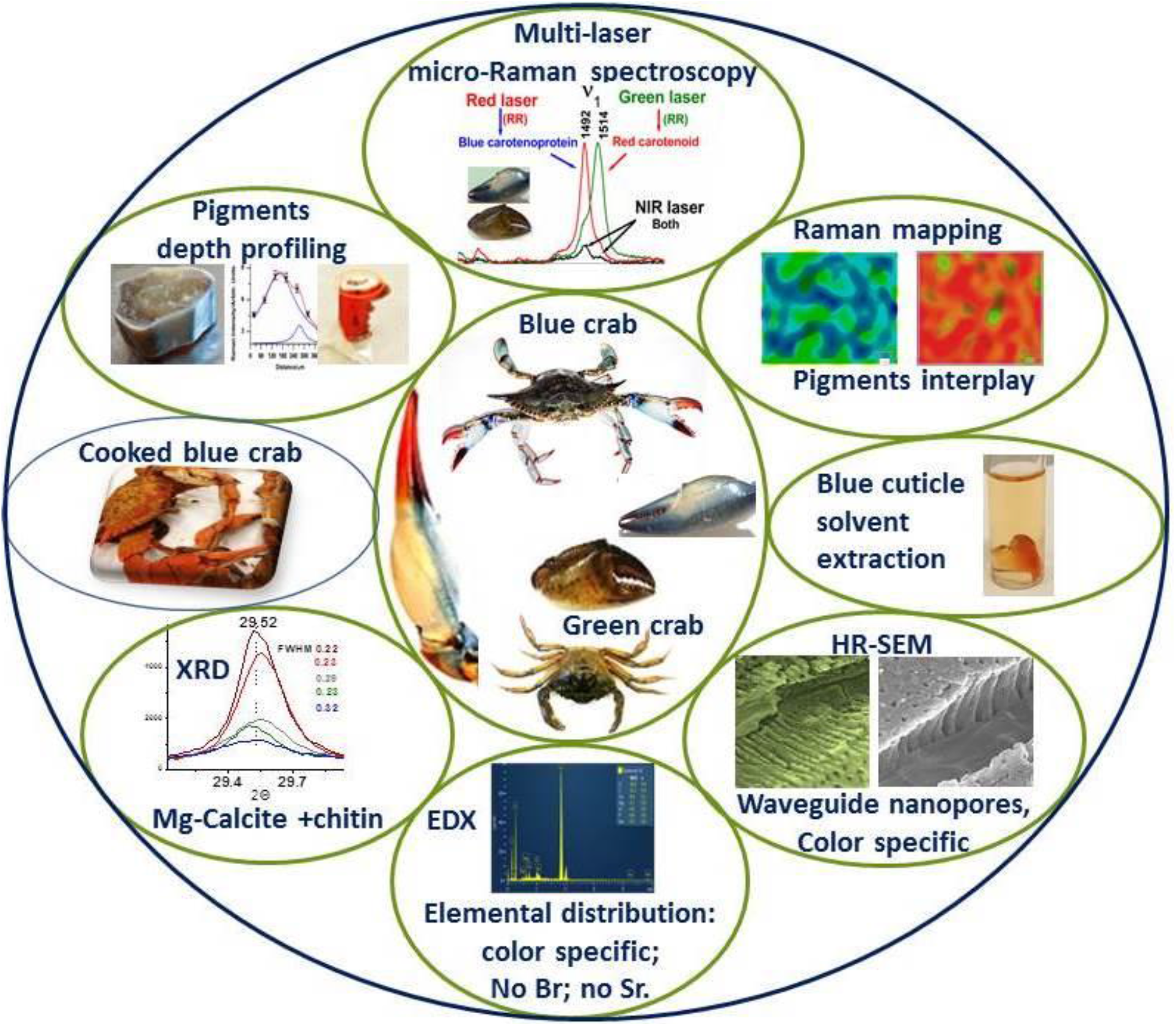
Summary display of the experimental approach comprising resonance and non-resonance Raman micro-spectroscopy, HR-SEM, EDX, XRD of blue, red, white and green exoskeleton of *C. sapidus* and *C. aestuarii* crabs in native form. Blue shells turning red when cooked or solvent extracted is molecularly illustrated by Raman spectroscopy.

### Multi-laser micro-Raman spectra of blue, red, green and white crab shells

Raman spectra collected from each crab shell color are strongly dependent on the laser excitation wavelength and highlights the free or ncb-ATX occurrence in the colored shells when proper excitation is applied (Supplementary Figure 2), Pigments overtones Raman analysis is given in Supplementary Figure 3. Comparative display of the carotenoid *v*_*1*_ *(C*=*C)* Raman band positions in spectra acquired from the four shell types with the three laser lines are showed in the Supplementary Figure 4. The **blue** exoskeleton fragments of *C. sapidus* cuticles under green laser excitation (532 nm) showed two Raman band components, one strong band assigned to the *v*_1_(C=C) skeletal stretching mode of ATX above 1500 cm^−1^ and a weaker band in the 1491-1493 cm^−1^ range, attributable to the same *v*_1_(C=C) mode in ncb-ATX in carotenoprotein. By contrary, full resonance Raman excitation of carotenoprotein with red laser (632.8 nm) line (which exactly fits crustacyanin extinction maximum^12^) readily revealed the RR mode arising from the ncb-ATX at 1492 cm^−1^ (Fig. 2) and denoted “blue band”. This signature is characteristic to the native carotenoprotein in crab shells and its presence with higher or lower intensity describes its relatively different abundance in various colored shells. Notably, its presence in green crab spectrum with lower intensity (Fig. 2), demonstrates the existence of ncb-ATX in blue carotenoprotein from *C. aestuarii*. In other words, blue pigment is present in green crab shell.

**Fig. 2.**
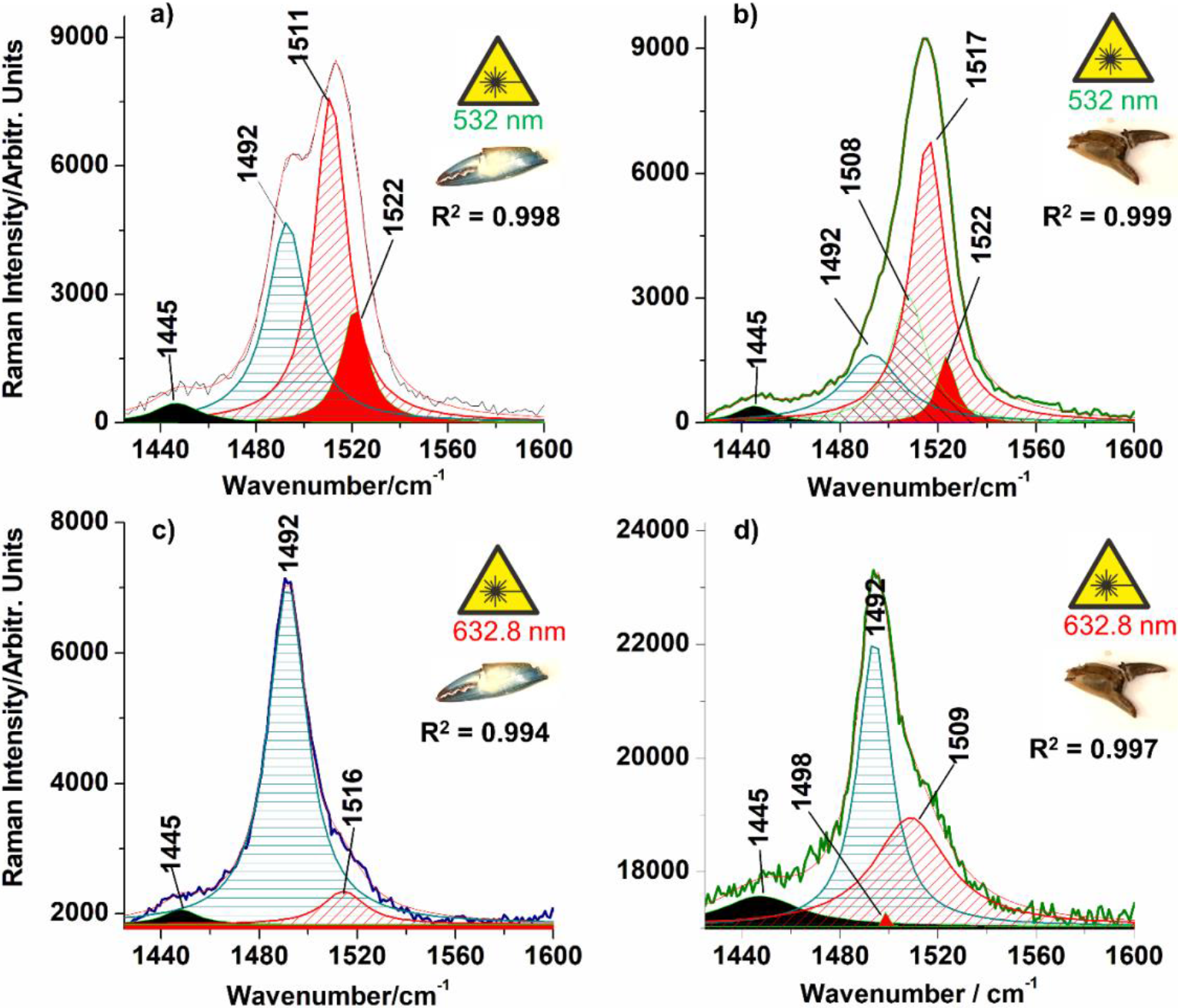
Multi-peak Lorentzian fit of raw RR spectra (1425-1600 cm^−1^ range) acquired from blue *Callinectes sapidus* claw shell (a, c), and green *Carcinus aestuarii* claw shell (b, d), using two laser lines, as indicated. Profile broadening via concentration was assimilated with pressure broadening, thus, Lorentzian fit was considered for multi-peaks deconvolution. Corresponding coefficients of determination (R^2^) are displayed in the figure. Note the effect of resonant excitation of pigments: deconvolutions of spectra excited with 532 nm featured stronger carotenoid modes above 1500 cm^−1^, while ncb-ATX mode at 1492 cm^−1^ is always dominant as “blue band” in spectra excited with the 632.8 nm line. Additional -CH_2_ mode (filled black band) at 1445 cm^−1^ along with other carotenoid species with minor overall contribution, are highlighted.

The greater wavelength laser lines (785, 830 and 1064 nm), which are neither resonant to ATX nor its protein complex, always featured a weak shoulder of free ATX band around 1514 cm^−1^ associated with the stronger mode of ncb-ATX at 1491 to 1492 cm^−1^ (Supplementary Fig. 2). All Raman bands recorded from the blue shell with 5 laser lines are listed in Table 1 and their assignments are provided according to our computational chemistry results and literature^24, 35^. We conducted theoretical calculation of the RR spectrum of non-covalently bond ATX (ncb-ATX) and compared it to its free counterpart, to correctly assign the experimentally observed Raman data. Computed Raman modes of isolated ATX featured theoretical *v*_1_ band at 1508 cm^−1^, arising from conjugated in-plane C=C stretching modes, while the calculations for ncb-ATX (ncb-ATX) involved in strong dipole-dipole interactions, which simulated non-covalent binding to the protein carrier, exhibited *v*_1_ mode red shifted to 1494 cm^−1^ (Supplementary Fig. 5). Our calculated *v*_1_ positions are similar to respective experimental modes of α-crustacyanin reported in older and more recent studies^12,15,17^, which indicated good simulation of Raman response of ATX bound in carotenoprotein. The second Raman band themed *v*_2_ assigned to the C-C stretching mode of carotenoid skeletal structure was found at 1154 cm^−1^, being non-sensitive to the non-covalent interaction.

**Table 1.** Experimental Raman data/cm^−1^ of blue *C. sapidus* claw cuticle obtained with different laser lines (532, 632.8, 785,830, 1064 nm) comparatively showed with the theoretical calculated Raman modes of ATX and ncb-ATX and their assignment.

| Raman experimental data/cm<sup>-1</sup><br>of blue *C. sapidus* cuticle |  |  |  |  | Calculated Raman<br>modes/cm<sup>-1</sup>:<br>free ATX, ncb-ATX | Vibrational assignment |
| --- | --- | --- | --- | --- | --- | --- |
| 532<br>nm | 632.8<br>nm | 785<br>nm | 830<br>nm | 1064<br>nm |  |  |
|  |  |  | 154 | 155 | - | Calcite (T) |
|  | 281 | 282 | 284 | 281 | - | Calcite (L) |
| 712 | 712 | 712 | 714 | 712 | - | $\nu_4(\text{CO}_3^{2-})$ in calcite |
| - | - | 880 | - | - | 875,882 | ATX $\delta(\text{CH}_2)$ |
| - | - | - | - | 892 | - | α-chitin |
| 953sh | 953sh | 953sh | - | 955 | 923,929 | ATX $\nu(\text{C-C}) + \rho(\text{CH}_2) + (\text{CH}_3)$,<br>α-chitin, $\nu(\text{PO}_4^{2-})$ |
| 963 | 963 | 963 | 963 | 963sh | 972sh,972sh | ATX $\nu_4(\text{C-C}) + \rho(\text{CH}_2) +$<br>$(\text{CH}_3)$, α-chitin, $\nu(\text{PO}_4^{2-})$ |
| 1004 | 1004 | 1003 | 1004 | 1004 | 992, 997 | $\nu_3$ ATX, ncb-ATX $\rho(\text{C-CH}_3)$,<br>Phe |
| - | - | - | - | 1038 | - | proteins (Phe) |
| 1086 | 1088 | 1086 | 1087 | 1086 | - | $\nu_1(\text{CO}_3^{2-})$ in calcite |
| 1154 | 1154 | 1154 | 1154 | 1154 | 1154, 1154 | ATX and ncb-ATX $\nu_2(\text{C-C})$, α-<br>chitin |
| 1190 | 1190 | 1197 | 1190 | 1190 | 1186,1189 | ATX, ncb-ATX $\nu(\text{C-C})$ |
| | 1205 | 1203 | 1206 | 1202 | - | α-chitin |
|  | 1234 | 1234 | 1234 | 1234 | - | proteins (amide III) |
| 1265 | 1264 | 1263 | 1263 | 1263 | 1262,1269 | ncb-ATX, ATX $\rho$(C-H), α-chitin |
| 1301 | 1305 | 1304 | 1303 |  | 1314, 1307 | ncb-ATX in crustacyanin |
| | | | | 1325 | - | α-chitin |
| 1385 | 1375 | 1374 | 1375 | 1375 | 1384,1384 | ATX, α-chitin |
| | | | | 1412 | - | α-chitin |
| 1444 | 1453 | 1442 | - | 1448 | 1444,1425 | ATX, ncb-ATX $\delta_{\text{asym}}(\text{CH}_{2,3})$, α-chitin |
| 1492 | 1492 | 1492 | 1492 | 1492 | absent,1494 | $\nu_1(\text{C}=\text{C})_{\text{out-of-plane}}$ in ncb-ATX |
| 1514 | - | 1514 | 1514 | 1514 | 1508, absent | ATX, $\nu_1(\text{C}=\text{C})_{\text{in-plane}}$ |
| | | 1517 | | 1517sh | absent, 1522 | ncb-ATX $\nu_1(\text{C}=\text{C})_{\text{in-plane}}$ |
| - | 1603 | 1604 | - | - | absent,1609 | ATX, ncb-ATX $\nu(\text{C}=\text{C})_{\text{out-of-plane}}$ + |
| | 1621 | 1619 | - | 1621 | - | Proteins (Trp, Tyr, Phe) + $\nu(\text{C}=\text{O})$, α-chitin |
| - | - | 1657 | - | 1655 | - | α-chitin, proteins (amide I), α-chitin |
| - | - | - | - | 1746 | - | Calcite ($2\nu_2$) |
| 2117 | 2117 | absent | absent | absent | - | $\nu_2 + \nu_4$ in ATX, ncb-ATX |
| 2158 | 2158 | absent | absent | absent | - | $\nu_2 + \nu_3$ in ATX,ncb-ATX |
| 2308 | 2308 | absent | absent | absent | - | $2\nu_2$ in ATX, ncb-ATX |
| absent | 2448 | absent | absent | absent | - | $\nu_1 + \nu_4$ in ncb-ATX |
| 2518 | absent | absent | absent | absent | - | $\nu_1 + \nu_3$ in ATX |
| absent | 2646 | absent | absent | absent | - | $\nu_1 + \nu_2$ in ncb-ATX |
| 2668 | absent | absent | absent | absent | - | $\nu_1 + \nu_2$ in ATX |
| 3028 | absent | absent | absent | absent | - | $2\nu_1$ in ATX |
| - | - | 2877 | - | 2878 | - | α-chitin $\nu(\text{CH}_{2,3})$ |
| - | - | 2937 | - | 2934 | - | α-chitin $\nu(\text{CH}_{2,3})$ |
| - | - | 2969 | - | 2965 | - | α-chitin $\nu(\text{CH}_{2,3})$ |

Abbreviations: ATX-astaxanthin, ncb-ATX - non-covalently bound astaxanthin, $\nu$ =stretching, $\rho$ =rocking, $\delta$ =bending, T, L -lattice modes in calcite.

Correlated experimental and theoretical findings clearly suggest that blue shell contains both free and ncb-ATX in carotenoprotein complex. Harmonics and linear combinations of the *v*_1_, *v*_2_, *v*_3_ and *v*_4_ modes of ATX or ncb-ATX in the high wavenumber range (2100 - 3100 cm^−1^) confirmed the existence of two pigments in the blue shell: they corresponded to ATX modes (Table 1) when the shells were excited with green laser line, and to ncb-ATX, under red laser excitation (Supplementary Fig. 3).

**The red and white** *C. sapidus* claw cuticles as well as the green ones of *C. aestuarii* have been similarly investigated, as comparatively shown in Supplementary Fig. 2 b-d. Excitation with 532 nm line revealed the presence of ATX, while the 632.8 nm line revealed both signal of ncb-ATX and free ATX in red and green shells. Weak and noisy Raman signal of pigments has been spuriously detected in white shells, too (Supplementary Fig. 2d). Thus, it appears that the presence of ncb-ATX in crustacyanin complex is not limited to blue shells of *C. sapidus* only. In other words, shells containing the complex are not necessarily blue, similar to findings on yellow lobsters^11^, which exhibited strong RR signal of yellow protein, and weak RR signal of ncb-ATX from crustacyanins.

Multi-peaks Lorentzian fit of the carotenoid *v*_1_ band in Raman spectra acquired from blue *C. sapidus* and green *C. aestuarii* claw shells (Fig. 2) provides accurate bands composition, revealing other, minor contributions to the main C=C band shape (R^2^ > 0.99). It supports the resonance excitation of free ATX with 532 nm line, and ncb-ATX in crustacyanin complex with 632.8 nm line. The difference in position of C=C modes above 1500 cm^−1^ can be attributed to stretching modes of conjugated C=C bonds of free carotenoids. The C=C mode arising from the ncb-ATX complex consistently appeared 1492 cm^−1^ (showed as blue shaded band in Fig. 2).

To further clarify the different colors of shells which contain the same pigments, we calculated the relative Raman intensity ratio (R = I_1492_ / I_1514_) of the two main modes contributing to the overall *v*_1_ Raman band, assigned to the ncb-ATX and ATX. The near-infrared 785 nm laser line was used for this purpose, as it provides normal Raman scattering only, thus avoiding resonance Raman contribution of either pigment. As the intensity of normal Raman scattering is directly proportional to the analyte concentration, for similar excitation conditions and collecting optics, the intensity ratio (R = I_1492_ / I_1514_), gives an estimation of relationship between the content of ncb-ATX and free ATX chromophores in each native shell color. The calculated values of the R ratios defined above are shown in the **Table 2** along with their standard errors. Data processing algorithm for the calculation of the R ratio values and the averaged corresponding spectra involved are shown in the Supplementary Fig. 6.

**Table 2.** Calculated Raman intensity ratios (R) of free- and non-covalently bonded ATX chromophore modes in blue, red and green cuticle, reflecting relative pigment content.

| Shell color type | Pigment Raman ratio: $I_{1492} / I_{1514}$ (mean ± S.E.) | number of measurements |
| --- | --- | --- |
| blue | $8.487 \pm 0.283$ | 9 |
| red | $0.757 \pm 0.301$ | 12 |
| green | $2.958 \pm 0.277$ | 10 |

S.E.- standard errors.

Note the highest value of R = 8.487 ± 0.283, for the blue color, almost 3 times higher than in the green shell, and more than 11 times higher than in the red shell, meaning that blue shells contain the most ncb-AXT chromophores relative to AXT. Due to the natural inhomogeneity of shell surface color, the considerable standard error values are not surprising. No carotenoid signal was recorded from white shells with non-resonant 785 nm excitation due to their low content. Summarizing, different colors of *C. sapidus* and *C. aestuarii* cuticle arise from the balance of co-existent orange and blue pigments. Surface distribution of pigments in crabs cuticle has been further mapped to support the standard errors of R values described above. StreamLine™ Raman imaging approach allowed for rapid generation of Raman images of lateral distribution of ATX and ncb-ATX signal on surface of blue and red *C. sapidus* and dark green *C. aestuarii* dorsal claw cuticle (Fig. 3). Attempt to do the same analysis for the red shells failed due to the much stronger RR signal + background of ATX, which completely covered any trace of the ncb-ATX complex contribution in the StreamLine™ data collection. The obtained Raman images confirmed the co-existence of both pigment types, their inhomogeneity distribution and their interplay.

**Fig. 3.**
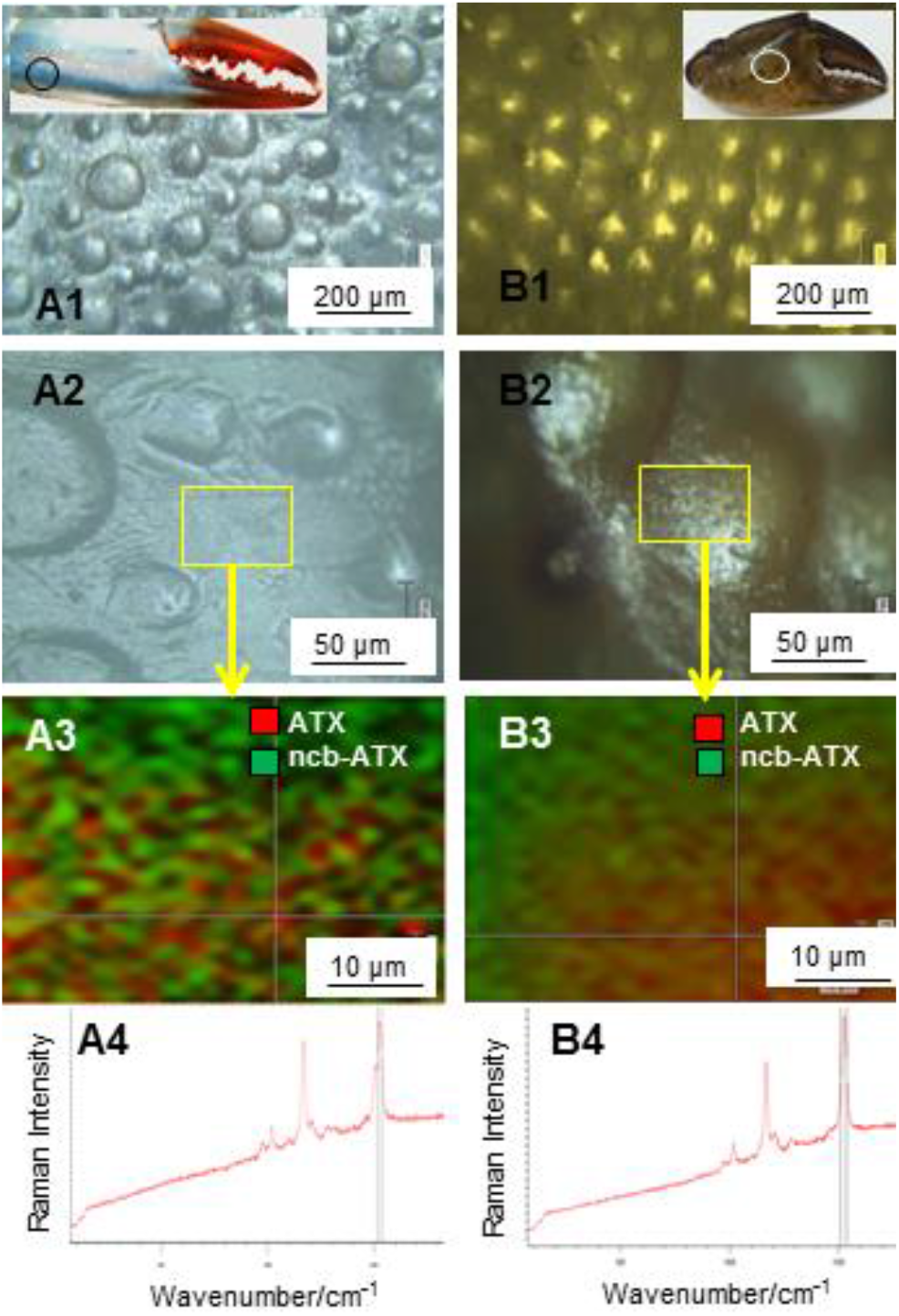
Mapping the Raman intensity distribution of the astaxanthin (ATX) *v*_1_(C=C) band at 1514 cm^−1^ and its non-covalently bound counterpart (ncb-ATX) at 1492 cm^−1^ from spectra collected from the blue *Callinectes sapidus* and green *Carcinus aestuarii* crab cuticle surface. Excitation: 532 nm. A1,2 and B1,2 show the light microscopy images taken via Raman microscope with 5x or 20x objective respectively, while A3 and B3 display the two pigments interplay over the mapped area highlighted in rectangle in A2 and B2. A4 and B4 show the rough Raman signal of the respective cuticle at the cross-hair of the maps. Note the spatial interplay of the two pigments contribution.

#### To further get insight into the shells properties, we investigated the pigments distribution in their cross-sections

By cleaving shells natural fracture planes were obtained and tracked by Raman micro-spectroscopy following normal-to-surface direction. Lateral step ranging from 20 to 60 μm was carefully chosen to comprise as many points as possible with relevant pigments signal, taking into account the thickness variability of the cuticle along the claws or palms. The ATX and ncb-ATX exhibited well-defined Raman signal and the intensity evolution of carotenoid *v*_1_ band was tracked, as showed in the Fig. 4.

**Fig. 4.**
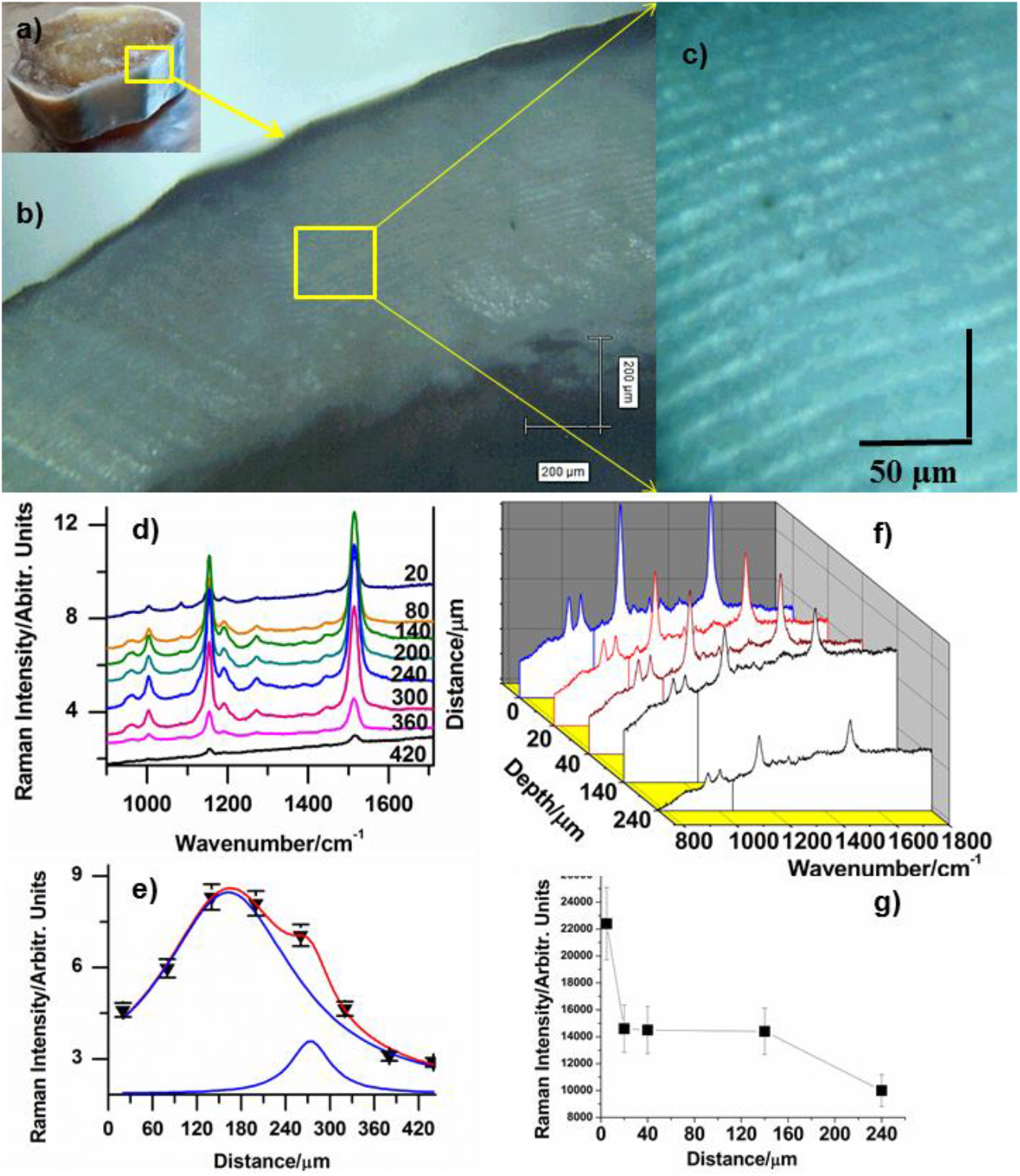
Pigments distribution in the cross section of blue cuticle (a) along with the normal-to-surface direction: b-c) micrographs taken via Raman microscope during measurements; d) RR spectra of ATX collected at various depths, from 20 to 420 μm as indicated on each spectrum; e) ATX Raman C=C mode (1514 cm^−1^) intensity distribution against depth; the data fit showed a Lorentzian profile with two components, peaking at 130 and 260 μm respectively; f) RR spectra of ncb-ATX taken from surface to 240 μm depth; and its main band (1492 cm^−1^) intensity variation along cross section g) showing the highest intensity in epicuticle. Error bars show the standard deviation.

The fit of collected data points revealed two distinct depths with high ATX content in blue shell, the first maximum being around 130 μm, and the second at the depth of about 260 μm (Fig. 4). Red and green cuticles exhibit different depth profile of ATX band intensity (Supplementary Fig. 7). Cross section of blue cuticle tracked with the 632.8 nm line revealed highest intensity of ncb-ATX in epicuticle, following a rapid decrease within 20 μm depth, than further slow decrease between 20-140 μm and very weak intensity up to 240 um depth (Fig. 4). Previous immunostaining approach to determine pigments distribution in spiny lobster *Palinurus cygnus* ^34^ located crustacyanins only at the depth of 7 to 8 μm beneath cuticle surface and in hypodermis of the cuticle.

### Blue and green shells turn red-orange when immersed in boiling water or after ethanol extraction, leading to the disappearance of the ncb-ATX resonance Raman band

Additionally, when blue, red and green claw shells were immersed in boiling water (100 °C) or ethanol immersion to probe pigments extraction, Raman analysis of same shell fragments showed that intact AXT-crustacyanin complex is prerequisite for maintenance of shell native color. Each treatment resulted in complete dissociation of the complex (Supplementary figure 8), leaving behind abundant free AXT, which gives intense orange color aspect to the shells. In addition, ethanol extraction appeared simple method of obtaining valuable carotenoid-enriched solution from the shells, which may further be used for carotenoid purification, while preserving the intact porous calcite morphology. (i) *C. sapidus* blue and red, and *C. aestuarii* green claw shells changed their native color into red-orange after few seconds of immersion into boiling water (Supplementary Fig. 8a, b). White couterparts of *C. sapidus* claw remained unchanged. Only ATX could be detected in the treated shells (Supplementary Fig. 8b), while the ncb-ATX RR band was absent. It results that crustacyanin complexes released ATX at boiling water temperature, followed by diminishing of darker color shade, while free AXT, which would not be affected by short exposure to high temperature, remained unaltered. (ii) Likewise, the blue *C. sapidus* native claw shell turned pinkish red after 14 days of carotenoid extraction in ethanol. The marker RR band of ncb-ATX was not detectable in shells after extraction, indicating total dissociation of the ATX-crustacyanin complex by ethanol. Furthermore, the additional RR analysis of extract solution showed that only free ATX may be retrieved from crab shells after exposure to ethanol, due to the dissociation from crustacyanin under the effect of polar protic solvent treatment.

#### Mineral composition of the crab claws

Fourier-transform Raman (FT-Raman) data obtained with the 1064 nm laser line excitation revealed the prominent symmetric stretching mode of carbonate *v*_1_(CO_3_^2−^)^39^ in calcite, ranging from 1085 to 1088 cm^−1^, the *v*_4_ *in plane* bending mode^35^ around 712 cm^−1^ and well-defined calcite lattice bands at 155 and 281 cm^−1^ in all cuticle color types (Supplementary Fig. 9). The intensity and full width at half maximum (FWHM) values of the^−^ *v*_1_(CO_3_^2−^) stretching mode indicated the highest calcite crystallinity in red dactyls, followed by green dactyls, white, and blue palm shells. Significant broadening of the *v*_1_ mode to the lower wavenumbers, paired with broadening of the *v*_4_ and lattice modes indicates the co-existence of amorphous^39^ CaCO_3_ in blue and white shells, while broadening of *v*_1_ to higher wavenumbers indicates small inclusions of Mg atoms in calcite crystalline lattice^36^. The area ratio r of the two Raman band components of calcite stretching mode in blue shell showed a value r = area_(1085)_/area_(1075)_ of 2.82 (Supplementary Fig. 9). Additionally, α-chitin^37,38^ could be identified according to its characteristic Raman fingerprint bands at 892, 955, 1059, 1111, 1147, 1202, 1263, 1325, 1375,1412, 1448, 1621, 1655 cm^−1^ as well as the CH_2,3_ stretching modes at 2878, 2934 and 2965 cm^−1^ (Supplementary Fig. 9, Table 1). Weak protein bands^39^ were observed at 1001, 1038, 1234 cm^−1−^, and possibly contributing to bands at 1621 and 1655 cm^−1^, (Table 1). Carbonates, α-chitin, proteins and carotenoids all account for the observed FT-Raman bands with variable relative intensity from one beamed point to another. Such relative distribution over the cuticle surface is illustrated in the Raman maps generated from the signal-to-baseline approach in fast StreamLine™ imaging of blue cuticle surface (Supplementary Figure 10) excited with the NIR line at 785 nm line.

The X-ray powder diffraction data confirm the presence of low Mg-calcite (the (1 0 4) peak at d=3.02, compared with 3.035 in pure calcite), according to results of Boßelmann et al.^40^ reported for *H. americanus* and *Cancer pagurus*. The highest Mg-calcite pattern was observed in red claw (even higher for its corresponding teeth) followed by the white, green and blue cuticle, as showed in the Fig. 5. The FWHM (showed in the insert of Fig. 5) varied from 0.23° in red and green, 0.29° in white to 0.32° 2θ in the blue cuticle. The dominant Mg-calcite of variable crystallinity is associated with small amounts of quartz (probably originated from benthic diatoms inhabiting the cuticle and/or spurious sand micro-grains) in some of the shell parts (especially in the red shell and claw teeth) observed at 26.67° 2θ, while a weak, broaden band centered at 19° 2θ was assigned to α-chitin^41,42^.

**Fig. 5.**
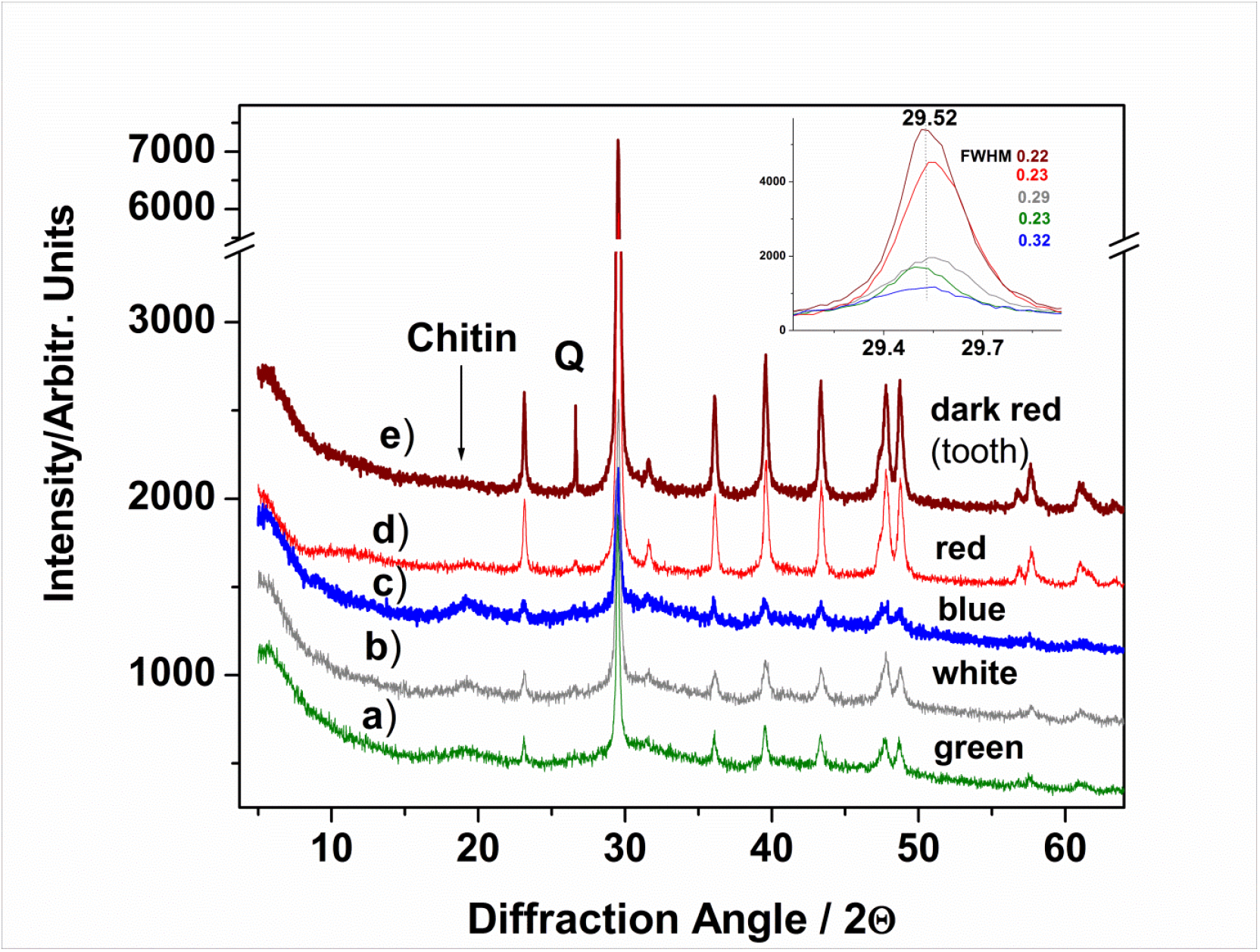
XRD diffractograms of the powdered crab cuticles as indicated, showing the Mg-calcite pattern. FWHM values of the main peak are inserted. Additional signal from crab tooth (e) is showed to highlight its higher crystallinity compared to the claw cuticles. Q-denotes quartz peak probably originated from benthic diatoms inhabiting the cuticle and/or spurious sand micro-grains (especially in the claw teeth). The weak band centered at 19° 2θ was assigned to α-chitin.

### Different crab color cuticles morphology showed systematic differences at ultrastructure level

An ordered, stacked mineral structure was observed, similar to those reported for other crab species^8,28–32^. A series of SEM images collected from fractured cuticles of blue shell is showed in the Fig. 6A. Comparative morphological details with the white and red cuticle of *C. sapidus* and green cuticle of *C. aestuarii* are given in the Fig. 6B. The averaged distances between pores and canals and their diameter for the white, blue, green and red cuticle are distinct, as showed in the Fig. 6C.

**Fig. 6.**
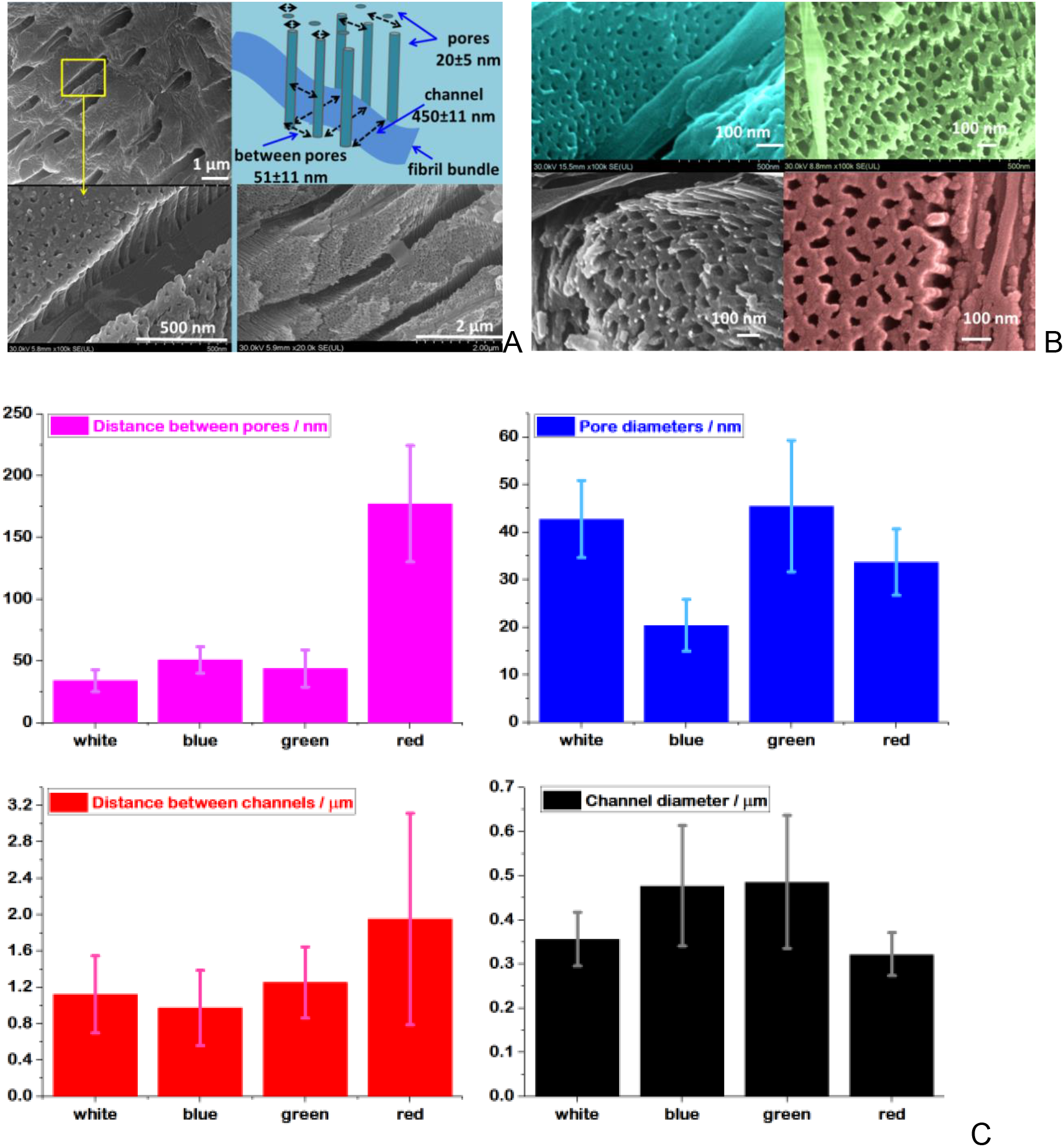
**A:** Representative SEM images collected from blue cuticle, highlighting the channels and the regular arrays of nano-pillars separated by pores in the ultrastructured channel walls. Schematic representation of the pores and channels arrangement is displayed in the top right corner. The channel width of about 450 nm is comparable with the blue light wavelength. **B**: Comparative SEM images from the four coloured shell, blue, green, white and red, showing nanoscale details of color-characteristic pores. **C:** The averaged distances between pores and canals and their diameter for the white, blue, green and red cuticle, as indicated on each graph. Error bars show the standard deviation.

The mineral layers are stacked quasi-parallel to the outer surface and their arrangement defines the epicuticles (detail of a detached epicuticle is showed in the Supplementary Fig. 11 for the green shell), the exocuticle, where the layers are thinner, and the endocuticle, where much thicker layers are observed. The mineral layers 3D nanoarchitecture is crossed by pore canals hosting the chitin-protein fibril bundles which follow a helical path normal to the cuticle surface. The canals determine the “macro-pores” which cross the successive mineral layers (Supplementary. Figure 11).

Such morphology explains the strong Raman scattering signal collected in back scattering configuration normal to cuticle surface, or weak signal when the collection direction is normal to the cross section of the cuticle. Furthermore, the spotted pattern of the pigments interplay from Fig. 3 might be consistent with the canals distribution, each canal being populated with chitin-proteins fibrils, thus strong Raman signal in spotted organic areas has been observed.

The pore canals exhibit complex arrangements, showing grating-like pattern of parallel mineral nanopillars (Fig. 6A), which also follows the helicoid path orientation. This structure, which has nanometer-scale arrangements at subwavelength optics, might be essential for selective light waveguiding through the canals, making the blue shell absorbance peaking between 550-650 nm, red between 400-500 nm, while the white one above 600 nm, which corresponds with the diffuse reflectance data previously reported^9, 33^. To further check any optical correlation, we measured a series of selected cuticle morphological features (Table 3). Each cuticle color revealed different width of pore canals and different distances between rows of pore canals, which in turn results in different grating constants responsible for manipulation of light. A schematic representation of these canals and pores acting as gratings is showed in the Supplementary Fig. 11.

**Table 3.**
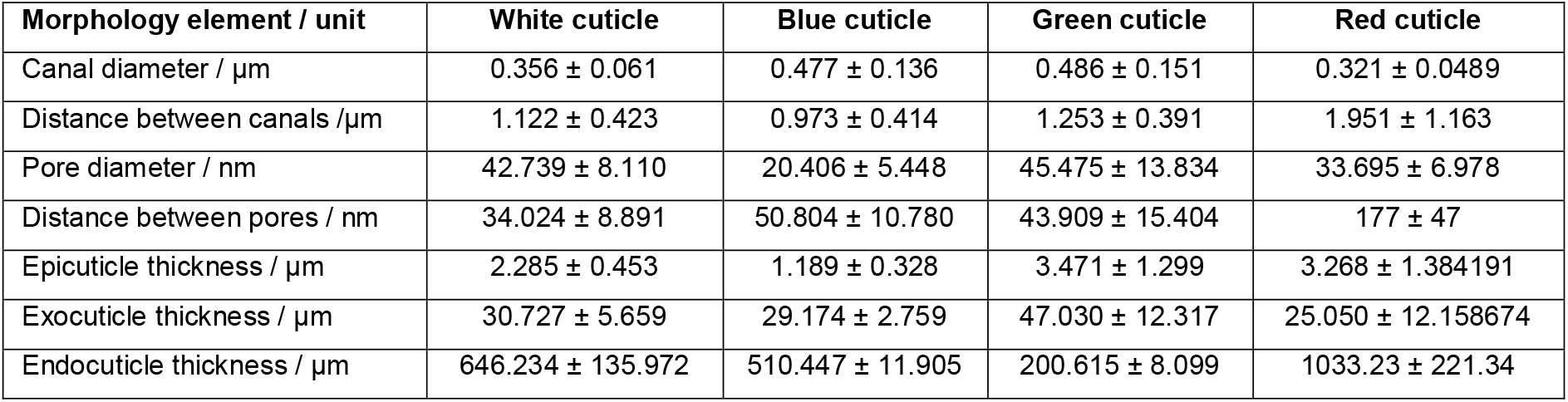
Comparative size of the measured crabs cuticle morphology elements. The data are listed as average values ± standard deviation.

Thus, the blue cuticle showed canals diameter (averaged values) of 477, green 486, red 321 and white 356 nm while the distance between canals was 977 nm in blue, the lowest among the other color cuticles, exceeding 1 μm (Table 3, Fig. 6C). The wall structure of canals showed the thinnest pores diameter in blue cuticle of about 20.4 nm, and the densest pore 3D-array. Combined with the thinnest epicuticle (Supplementary Fig. 12 and 13), the blue shell appeared morphologically distinct.

The finding overturns our understanding of the valuable 3d-nanomaterial with Bouligand patterns from blue crab shells, simultaneously exhibiting blue, red and white color.

Semi-quantitative elemental composition revealed by scanning electron microscopy and energy dispersive X-ray spectroscopy (SEM-EDX) supports the shell color distinct feature. Relative mass fractions (wt%) for the 5 main elements (Ca, C, O, Mg and P) were obtained from 2 to 4 scans (data plotted from individual column colors, Supplementary Fig. 13) corresponding to epicuticle, exocuticle and endocuticle of *C. sapidus* blue, red and white and *C. aestuarii* green cuticle, respectively (Supplementary Fig. 13). Semi-quantitative EDX elemental analysis (Supplementary Table S1) confirmed the FT-Raman and XRD conclusions of low Mg-calcite, by consistently revealing Mg in all shell color types and cuticle layers alongside Ca, C and O as main elements and trace of P, K, Na, and spurious S and Cl. Furthermore, detection of P in all shells, correlated with the Raman band at 963 cm^−1^ containing contribution of PO_4_ ^2−^ mode (Table 1), could indicate the presence of trace amount of carbonated hydroxyapatite, which is known to occur in crustacean cuticles^46,47^. However, this supposition is not supported by the XRD data showing that any phosphate mineral was under detection limit in the calcite matrix. Overall, the blue cuticle showed completely distinct composition, revealing the lowest Ca level in epicuticle, further increasing in exocuticle and showing an unusually high amount (up to 71.6 wt%) in endocuticle; lowest C, O and Mg level in endocuticle and highest P content in exocuticle. Such feature, could suggest distinct protective role of endocuticle as the thickest load-bearing layer. In line scan of elemental composition, EDX data constantly showed higher content of C associated to corresponding lower content of Ca, Mg and P, particularly in epicuticle, which mostly contains amorphous minerals and waxy lipoproteins ^43^.

## Discussion

Red ATX and its blue protein complex presence not only in blue, but in red or green crab cuticle is a proof of the complex mechanisms underlying the crustacean colors. Since the previous studies targeted mostly lobsters (*H. americanus* and *H.* gamarus)^10–19^, little is known about the occurrence and localization of astaxanthin-crustacyanin complexes in various crustacean species. Nevertheless, a genetic study conducted to prospect occurrence of genes that encode crustacyanin subunits^34^ established that crustacyanin is a lineage-specific adaptation developed by common ancestors of certain crustacean groups, however, the genome of *C. sapidus* was not considered.

Correlating the results, the pigments identity, their interplay and distribution in blue crab cuticle is clearly related to its biomineralization, which is different in blue, red white or green cuticles. Although the variability is high, as reflected by the standard deviation of data, the dimensions of the morphological micro- and nano canals and pores organized in regular 3D-nanoarchitecture clearly influence the selective light absorption, transmission, reflection, scattering or diffraction phenomena, thus, contributing to the final color appearance. The chemical composition and morphology showed well correlated Raman data with the XRD and SEM-EDX results, which revealed that the crab cuticle color is not governed by the pigments chemistry only, but also by their distribution in their specific 3D nano-architecture, which can be considered as a tridimensional supergrating.

Combined Raman spectroscopy approach including the proof of pigment identity by analyzing Raman overtones and linear combinations, although never applied in earlier Raman studies^12,15,17^ on carotenoids or carotenoproteins, seemed effective for native exoskeleton pigments analysis. For larger scale approach, Raman tools seemed effective for wasted materials assessment for blue bioeconomy purpose.

Although the cuticles exhibit low transparency^9,33^ for visible range, certain light wavelengths can reach, propagating through the channels and pores or scattering/diffracting on their typical 3D-nanoarchitecture. Thus, the blue shell showed a series of individual ultrastructural characteristics of pores, canals and nanopillars network which differentiate from the other shell colors. The aragonite polymorph reported in *C. sapidus* exoskeleton harvested on Aegean coast^33^, as well as the presence of Sr and Br to influence the crystalline structure and color^33^ is not sustained by the present data on blue crab harvested on Adriatic Sea coast.

Although the complex optical effects are difficult to be simultaneously assessed taking into account the pigments localization and weak light transmission in conjunction with the refraction, scattering, absorption, or diffraction and interference phenomena in the 3D super-array, their co-existence is supported by the physical-chemical and morphological characteristics. For example, chitin, a transparent polymer for the visible wavelength can modify the weak light transmission via refraction and scattering along the canals hosting the fibril bundles. The circular polarized light reflection which has been observed by Michelson in certain insects exoskeleton a century ago and supposed to be due to a “screw” structure (later named Bouligand pattern) could add additional contributions to the overall properties^53–55^ and color appearance not only for humans, but also for crustaceans vision. Other proteins can contribute with their specific absorption and/or fluorescence emission along the canals confined between cladding nanopillars walls network, while the nanopillars themselves defining the canals walls structure can act as 3D diffraction grating whose operability in certain diffraction order still has to be understood.

The highly ordered porous calcite 3D-supergrating, specific to each crab cuticle and gradually populated with balanced pigments appeared more complex than the extensively investigated structural colors in insects. In the latter, the optical properties of their cuticles stem from highly ordered Bouligand pattern comprising line gratings and/or vanes of their wings structure, with helicoid stacking of chitinous fiber layers^44,45^. *C. sapidus* cuticle may inspire further biomimetics avenues, since there are no clear-cut borders between blue, red and white shells. Since the crab cuticle can manipulate light with is combined chemical structure and morphology, light absorption and propagation in wavelength dependent manner could warrants attractive applications of the porous magnesian calcite supergratings. In the light of present findings, crab shells could potentially trigger the starting point of new, effective, advanced materials that could serve as porous model for preventing infections spreading^52^, control molecular solutions loading and releasing, bacterial therapies or develop new, effective, porous biostimulants for soils remediation and agricultural strategies, in line with the most innovative blue bioeconomy approach.

## Conclusions

Non-destructive multi-laser micro-Raman spectroscopy and imaging revealed that native claw shells in the Atlantic blue crab *C. sapidus* from invaded Adriatic Sea contain both the aggregated and the non-covalently bound astaxanthin in carotenoproteins. Further investigation of red and white *C. sapidus* and green *C. aestuarii* claw shells revealed that the two pigments are ubiquitously present in investigated shells, even in seemingly unpigmented white shells, albeit to much lesser extent. Thus, non-covalently bound astaxanthin in crustacyanins is not the only factor responsible for dark blueish and purplish color, but rather participates in formation of apparent color alongside free carotenoids and the 3D nanostructured regular array of Mg-calcite on organic scaffold. The pigments were found to co-localize over the cuticle surface and in its cross-section, with highest distribution in exocuticle layer. *C. sapidus* and *C. aestuarii* also may have a structural color component, derived from the 3D-nanostructured regular array of Mg-calcite on organic scaffold.

## Materials and methods

### Preparation of crab shells

Atlantic blue crabs (*C. sapidus*) and Mediterranean green crabs (*C. aestuarii*) were caught by gillnets and traps in Parila lagoon (southeast Adriatic Sea, Croatia)^2,6^. Crabs were brought to laboratory and frozen at −20 °C until needed. Before analysis, the crabs were thawed and their claws were dissected and cleaned from soft tissue that was inside using scissors, tweezers and tap water. Shells were then thoroughly washed in cold running tap water. Claw cuticles of 10 individuals, mainly sexually mature males and females, but also immature specimens differing in color shades were further cut into smaller pieces to fit the size requirements for respective analysis technique. Sample pieces of cuticle for analysis with each of the techniques described below were consistently taken from mature specimens featuring the most intense coloration of the claw palm. Comparative assessment has been conducted for their corresponding immature specimens with blue and white morphological areas. Red and green samples were taken from claw dactyls, as illustrated in several samples depicted in the Supplementary Fig. 1.

### Theoretical calculation

Geometry optimizations and DFT (Density functional theory) frequency calculations of the isolated astaxanthin molecule and astaxanthin near water molecules were performed with the Gaussian 09 software package^46^, at B3LYP/6-311G(d,p) level of theory^46–50^. The solvent effects have been considered by using the implicit SMD model^51^.

### Multi-laser micro-Raman spectroscopy analysis

Micro-Raman analysis was conducted on a Renishaw InVia confocal Raman microscope (Renishaw, UK). Four laser lines were used on this device: 532 nm excitation was provided by a Cobolt diode pumped solid state (DPSS), air cooled laser, He-Ne- laser provided the 632.8 nm excitation, and Infrared excitation was provided by two diode lasers emitting at 785 and 830 nm. Spectral resolution was 0.5 cm^−1^, and WiRE™ 3.4 software (Renishaw, United Kingdom) was used for operating the spectrometer and acquisition of spectra. The spectra were acquired either in single-point scan, or in StreamLine imaging mode (described later), from samples of cuticle mounted on microscope slides on a mobile XYZ HSES (high-speed encoded) stage.

Bruker Equinox 55 FT-IR spectrometer with integrated FRA 106/S IR Raman module was used for non-resonant Fourier transform excitation of crab shells. A Nd:YAG laser emitting at 1064 nm with 350 mW laser power was used for excitation, and signal was detected by a liquid nitrogen cooled Ge detector with spectral resolution of 2 cm^−1^.

The acquisition conditions, i.e. integration time, number of acquisitions per spectrum, laser power and objective magnification, varied according to specific sample surface characteristics (as discussed in Results section). The inhomogeneity of claw surface, both in relief and pigment content, made it challenging to obtain good focus and satisfactory signal-to-noise ratio, without exceeding the instrument count limit. Spectra acquired with 532 and 632.8 nm excitation lines featured incremental fluorescence background typical for carotenoid pigments while spectra acquired with 785, 830 and 1064 nm excitation featured low or negligible background level, due to greater laser wavelength which does not cause fluorescing effect of organic molecules.

Spectra of liquid sample, I.e. the astaxanthin-enriched ethanol extract, were acquired on a DeltaNu compact, dispersive Raman spectrometer (Intevac, United States). The 532 nm excitation line was used, and spectra were recorded with 8 cm^−1^ spectral resolution from samples prepared in 1-ml vials, using the NuSpec software (Intevac, United States).

### Streamline Raman imaging

Raman images were produced on surfaces of blue and red *C. sapidus* and green *C. aestuarii* claw shells using the StreamLine™ image acquisition facility of the Renishaw instrument. Both the 532 and 785 nm excitation lines were used separately, paired with 20x objective (NA 0.35). Firstly, a rectangular area to be analysed, 90 × 90 μm in size, was defined on the live video image. This area resulted in 9000 spectral acquisition points. The Raman image was acquired under a continuous laser beam by automatic movement of the stage to achieve 1 s exposure time per image pixel. The *v*_1_ mode of free ATX, centred around 1514 cm^−1^ and the *v*_1_ mode of ncb-ATX, centred around 1492 cm^−1^ were selected for creating separate images under the „signal to baseline” criterion. The images were then overlaid to produce composite pigment distribution maps.

### Spectral data processing

Raman and XRD spectra were processed in Origin 6.1 software (OriginLab, United States). Background subtraction was applied when needed, by creating a user-defined, automatic, 10 points baseline function and iteratively modifying it using the subtraction facility of the Origin software. Lorentzian fit multi-peak was applied for the Raman spectra in the 1425 to 1600 cm^−1^ range, estimating the initial half width and obtaining the peaks components position, area and center and yielding the coefficient of determination R^2^ higher than 0.99. Calculation of intensity ratio (R) of ncb-ATX band to ATX was done after background subtraction from spectra acquired from different shell color types with 785 nm excitation and subsequently comparing the intensity of the mode at 1492 cm^−1^ (ncb-ATX) and the mode at 1514 cm^−1^ (ATX) as R= I_1492_ / I_1514_ (Supplementary Fig. 6). Normalization of spectra was done by dividing all y data of spectral datasets by the intensity value of the desired band.

### Additional shells treatment

i. **High temperature denaturation of astaxanthin-crustacyanin complex**. Blue, red and white pieces of *C. sapidus* claw cuticle and green pieces of *C. aestuarii* claw cuticle were placed into Petri plate containing boiling water. Water was heated on a magnetic hot plate magnetic stirrer to 100 °C, and pieces of shells were kept in boiling water until their color changed to orange. Shells were then taken out of water using tweezers, dried on with paper towels, and subsequently analysed with the Raman microscope. Raman spectra were acquired from treated shells using the 532, 632.8, 785 and 1064 nm laser lines, and spectra were compared to those acquired from native blue and green shells.
ii. **Crude ethanol extraction of shell pigments.** Pieces of blue shells from *C. sapidus* claw were placed in vials containing 4 ml of ethanol. Raman analysis of both shells and the supernatant was conducted 14 days after the beginning of extraction. Shells and carotenoid microcrystals were analyzed with the Renishaw Raman microscope. The astaxanthin-enriched extract was analyzed in liquid state with the DeltaNu Advantage 532 Raman spectrometer.

### X-ray powder diffraction

Shell fragments were finely ground in an agate mortar and their mineral composition was verified by X-ray powder diffraction (XRPD), using a Bruker D8 Advance diffractometer in Bragg-Brentano geometry, with a Cu tube with λ_Kα_ = 0.15418 nm, Ni filter and a LynxEye detector. Corundum (NIST SRM1976a) is used as an internal standard. The data were collected on a 5 – 64° 2θ interval at a 0.02° 2θ step, measuring each step for 0.5 seconds. The identification of mineral phases was performed with the Diffrac.Eva 2.1 software (Bruker AXS) using the PDF2 (2012) database.

### Scanning electron microscopy and energy-dispersive x-ray spectroscopy

SEM imaging and EDX measurements were accomplished with a SU8230 Hitachi ultra-high resolution cold-field emission scanning electron microscope. The instrument allows for a combination of topographical and compositional information at a feature resolution of up to 1 nm in optimal conditions. Pieces of claw shells were dried in oven at 40 °C and kept in a container with CaCl_2_ bottom layer to avoid atmospheric water absorption. Before analysis, samples were adherently placed on Hitachi stub SEM holders (aluminium holder with M4 threads covered with carbon discs of 3 mm thickness). A Quorum Q150T sputtering sample coater capable of gold sputtering of controlled thickness for high resolution imaging, and evaporating carbon for EDX analysis was employed. Gold coating thickness was 10 nm (density 19.32 g/m^3^) at a rate of 14 nm/min. An Oxford energy-dispersive x-ray module (Oxford, UK) was used for elemental analysis of shells.

## Acknowledgement

The research leading to these results has received funding from the European Union Seventh Framework Programme (FP7 2007-2013) under grant agreement nr. 291823 Marie Curie FP7-PEOPLE-2011-COFUND (The new International Fellowship Mobility Programme for Experienced Researchers in Croatia - NEWFELPRO). This *paper* has been *prepared* as an additional part of the project “*Environmental aquaculture and seafood monitoring in South-Adriatic coast (Croatia) using Raman spectroscopy techniques and SERS-based sensors (acronym JADRANSERS)*” which has received funding through NEWFELPRO project under grant agreement nr. 5. This work was carried out on the Babeş-Bolyai University Research Infrastructure, financed by the Romanian Government through the programme PN II—Capacities - project “Integrated Network for Interdisciplinary Research (INIR)”, with direct public link to ERRIS: https://erris.gov.ro/Institute-of-Physics-IOAN-UR. We thank Professor Simion Astilean from Babes-Bolyai University for useful discussions and comments on the manuscript and particularly, on the light interactions with supergratings. Mr. Stanko from *Parilla Lagoon*, Croatia is highly acknowledged for prompt crabs capture whenever it was needed.

## Author contributions

F.N. and S.C.P. designed the research, conducted Raman spectroscopy analyses, collected, processed and analyzed data and wrote the manuscript, B.G. established the sampling area, selected the crabs specimens, assessed their biological status, and raised attention on possible specific pigments accumulation in wasted crab shells, specific for their invasive spreading area. V.C. produced the theoretical DFT calculations, L.B-T and M.S. conducted SEM-EDX data acquisition and interpretation, T.T. conducted the mineralogy assessment of the crab shells and recorded and analyzed the XRD data. All authors contributed to the data analysis, interpretation, manuscript writing and approved its final version.

## Competing interests

The authors declare no competing interests.

Corresponding author: Correspondence to Simona Cinta Pinzaru.

## Data availability statement

The data that support the findings of this study are available within the manuscript, its supplementary information and from the corresponding author upon reasonable request.

