## Supplementary Information for "Color-specific porosity in double pigmented natural 3d-nanoarchitectures of blue crab shell"

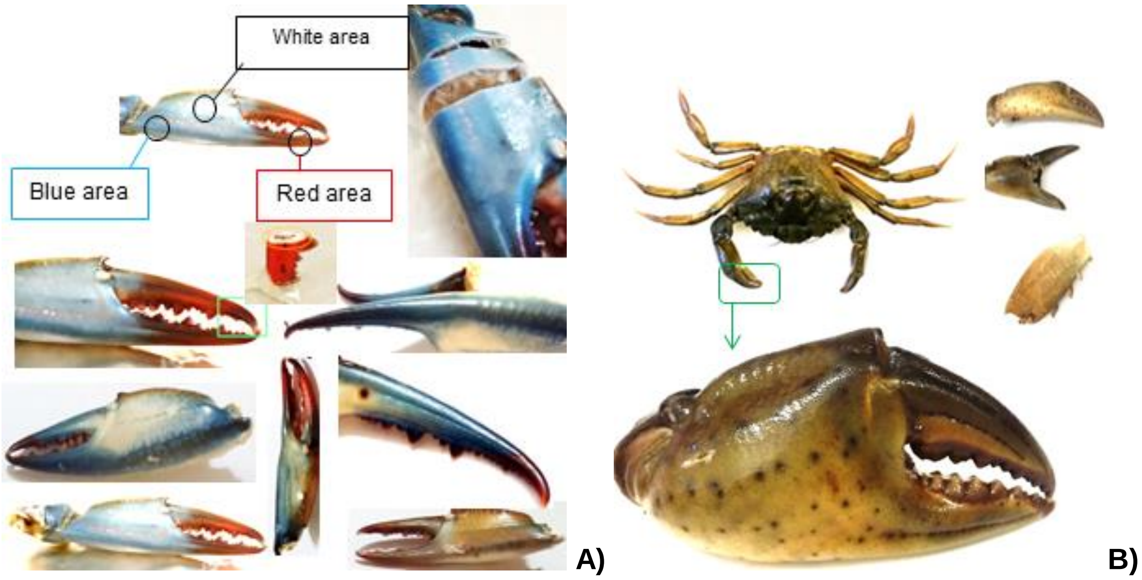


**Supplementary Figure 1. Photographs of the studied crab shells showing the diversity of color hues and their patterns. Morphological regions from which samples were taken are indicated as blue, red and white areas on *Callinectes sapidus* (the Atlantic blue crab) claws (A), and green area on *Carcinus aestuarii* (the Mediterranean green crab) claws (B).**

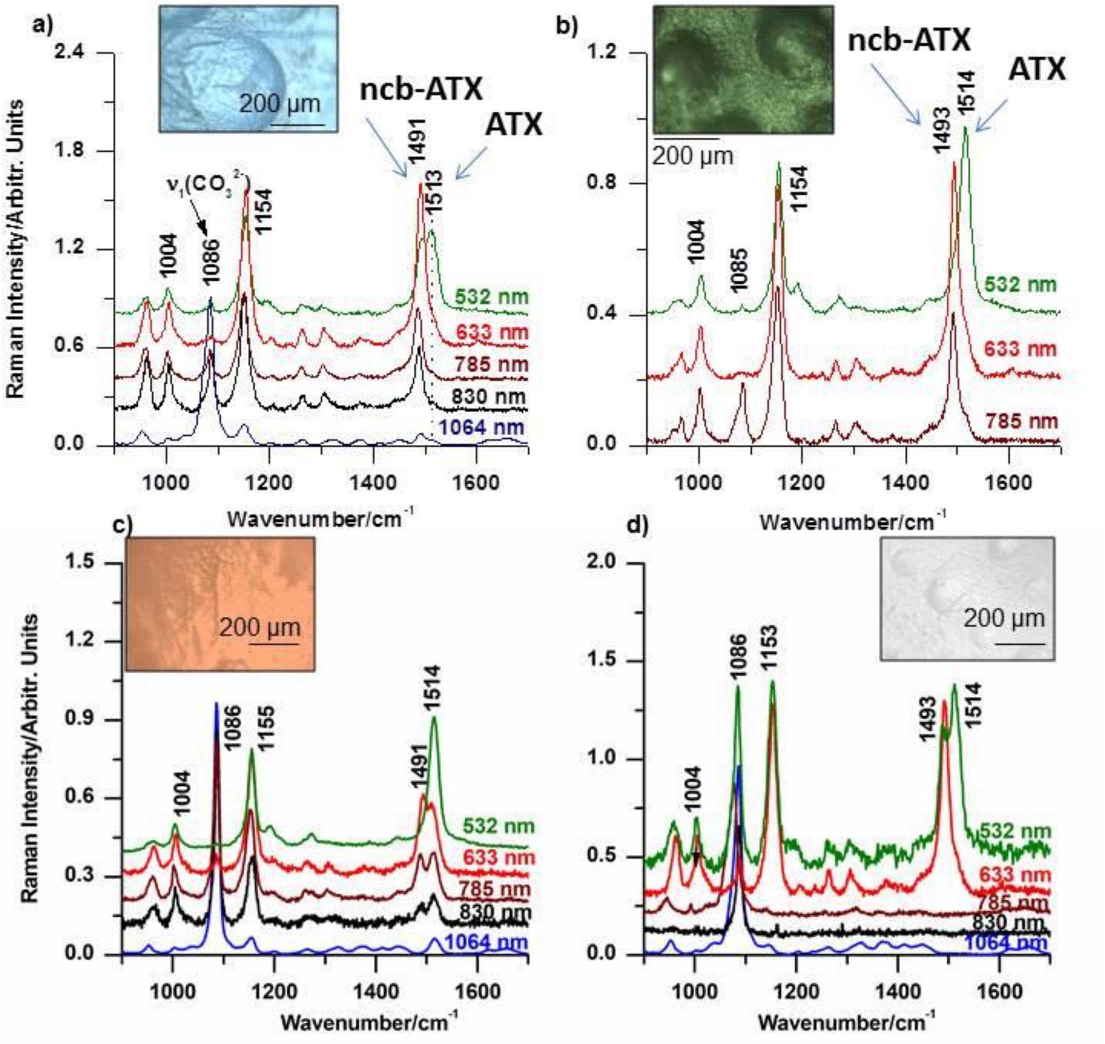


**Supplementary Figure 2, Multi-laser micro-Raman spectra acquired from a) blue, c) red, d) white *Callinectes sapidus* and b) *Carcinus aestuarii* claw shells, showing the co-existence of the free- and non-covalently bonded ATX in all the shell color types, respectively. Main carotenoid bands along with the $CO_3^{2-}$ mode (carbonate stretching) are labeled. Spectra were normalized and background subtracted. Corresponding micrographs (insets) taken from the shell surface under 200x magnification, highlight the intricate morphology and inhomogeneous color intensity. Laser excitation line is given on each spectrum.**

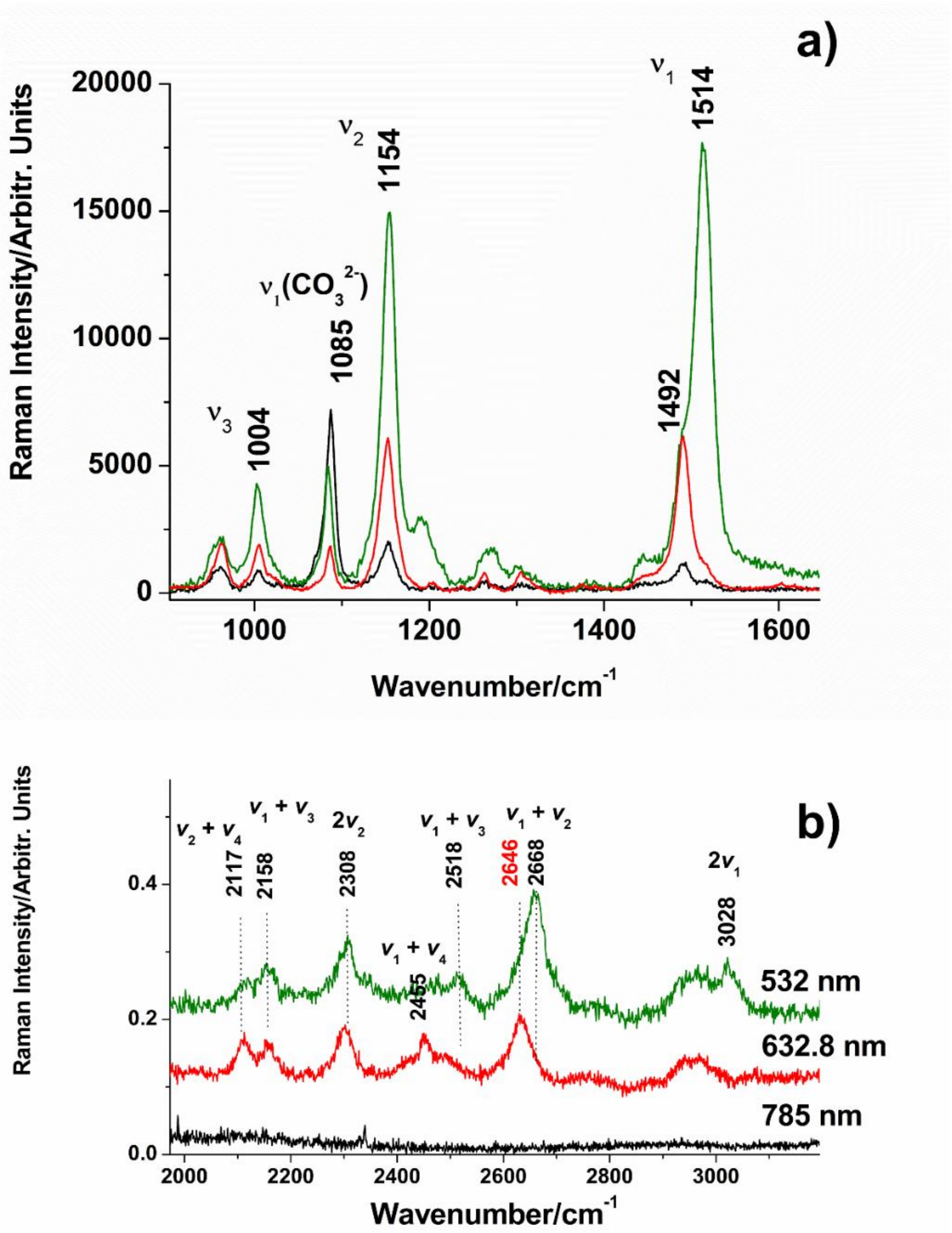


**Supplementary Fig. 3** Blue cuticle resonance Raman spectra of the free ATX excited with 532 nm (green spectrum) or ncb-ATX excited with 632.8 nm (red spectrum) in the fingerprint range a) and the corresponding overtones and linear combinations in the 2000-3200 cm$^{-1}$ range (b). Note the absence of these spectral features in the non-resonantly excited cuticle signal with 785 nm from b). Color codes are similar in a) and b).

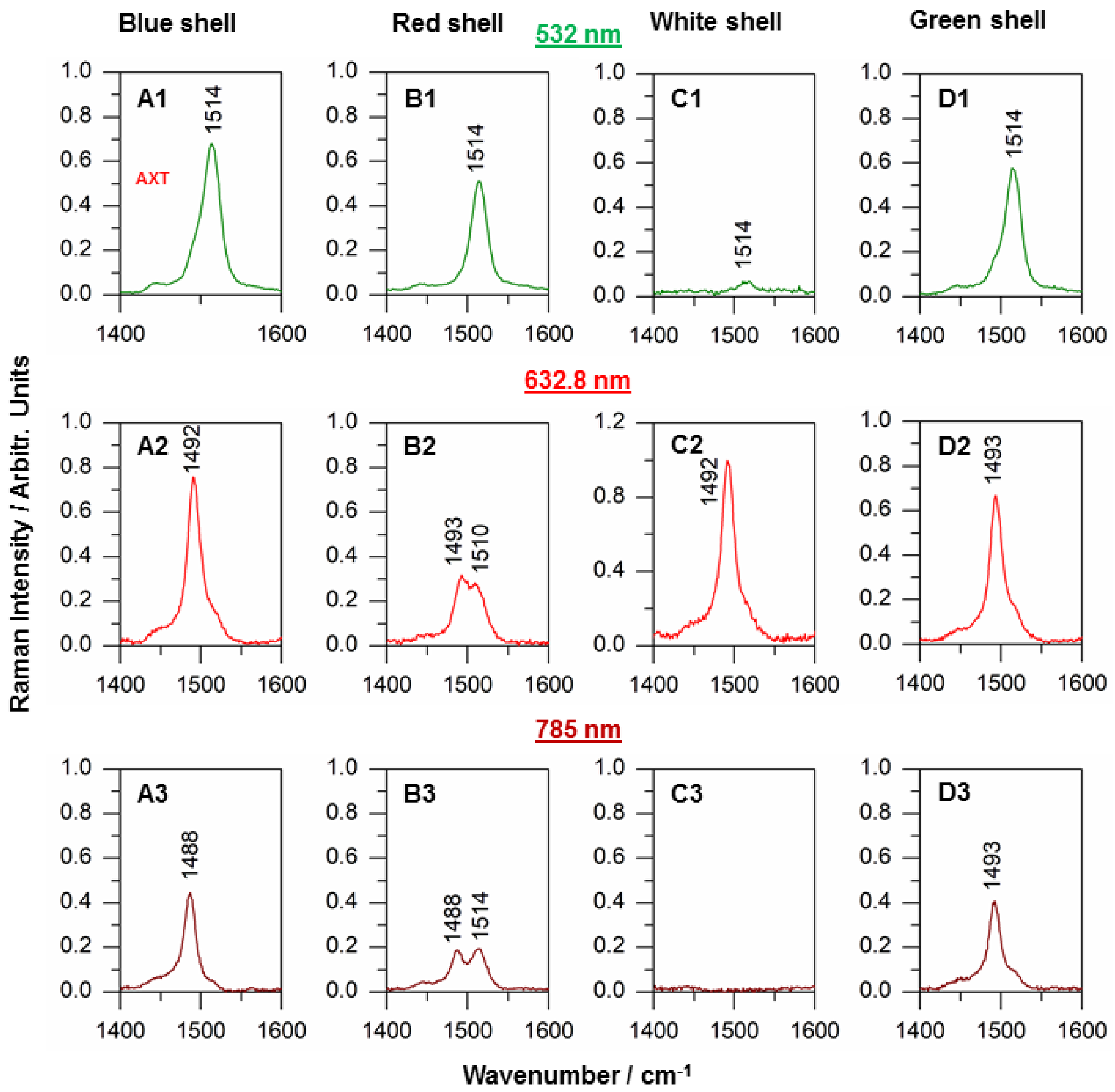


**Supplementary Fig. 4** Comparative display of the carotenoid $v_1$ Raman band positions in spectra acquired from the four shell types (vertically) with the three laser lines (horizontally), as indicated. Spectra are background-subtracted and normalized to the strongest signal. It can be clearly observed that all shell types contain both free- and non-covalently bonded astaxanthin (ncb-ATX), according to the bands at 1514 $cm^{-1}$ revealed by 532 nm laser line in each shell (top row), and around 1492-1493 $cm^{-1}$ revealed by 632.8 nm line (middle row). The 785 nm laser line is non-resonant to either ATX or ncb-ATX, thus, it excites the normal Raman scattering of both, coexistent pigments (bottom panels).

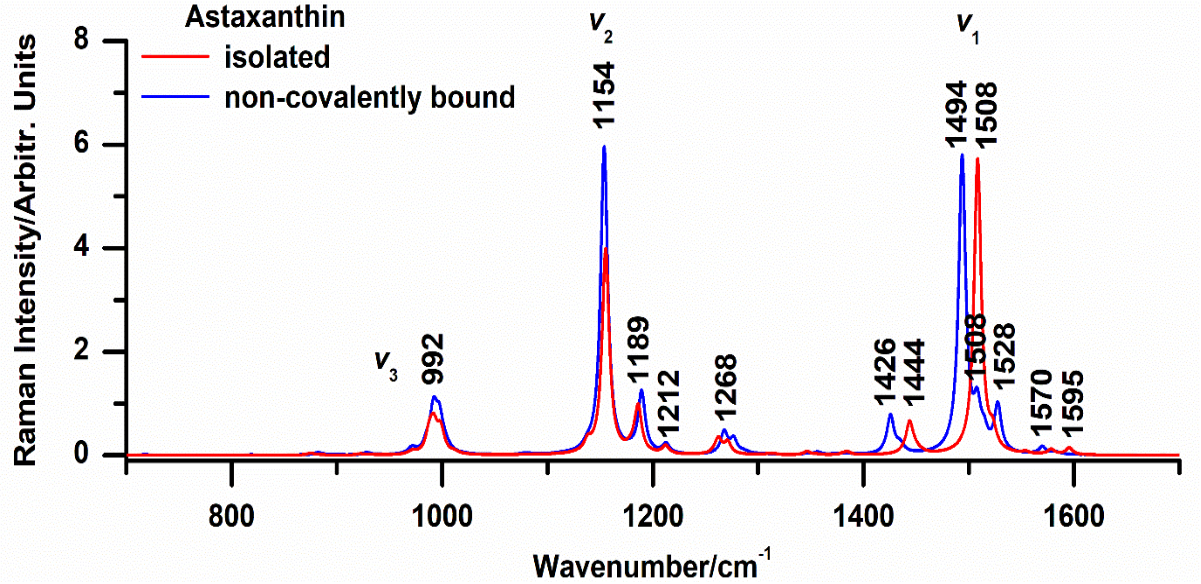


**Supplementary Fig. 5. Theoretical DFT calculated Raman spectra of isolated (gas-phase) astaxanthin (red line), and non-covalently bound astaxanthin in aqueous solution (blue line).**

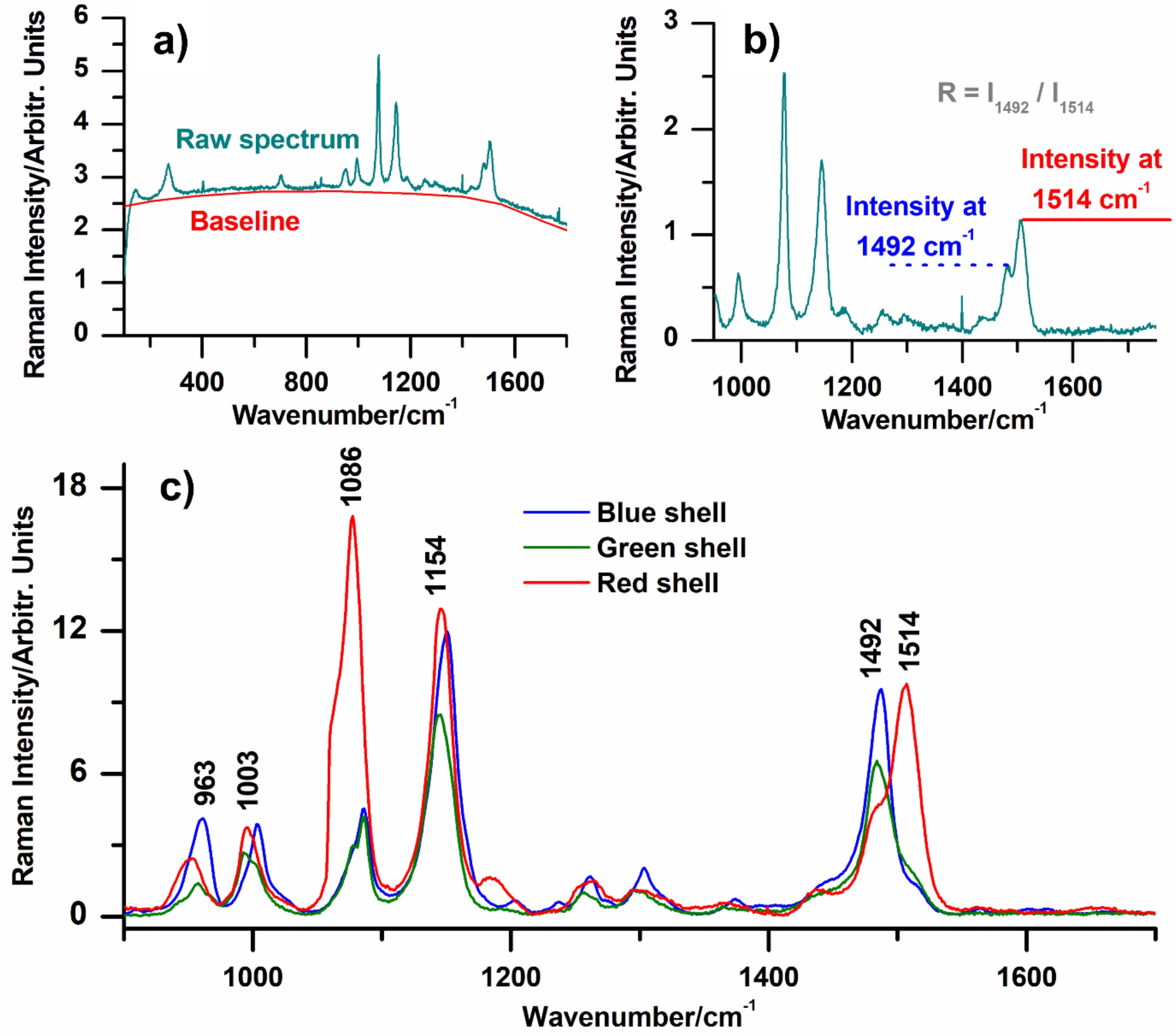


**Supplementary Fig. 6** Procedure of calculating pigment mode ratios using Raman spectra acquired from blue, red and green claw shells non-resonantly excited with the 785 nm line. a) baseline is removed from spectra, b) intensity of ncb-ATX mode around 1492 $cm^{-1}$ and astaxanthin mode at 1514 $cm^{-1}$ is measured. The ratio is calculated as $R = I_{1492} / I_{1514}$. c) averaged, baseline subtracted spectra normalized to $CO_3^{2-}$ mode around 1086 $cm^{-1}$, from the three shell color types, as indicated. Note that the crustacyanin contribution to $v_1$ is the highest in blue shells, exhibiting the highest R value, followed by green shells, while the red cuticle exhibits stronger contribution from free ATX.

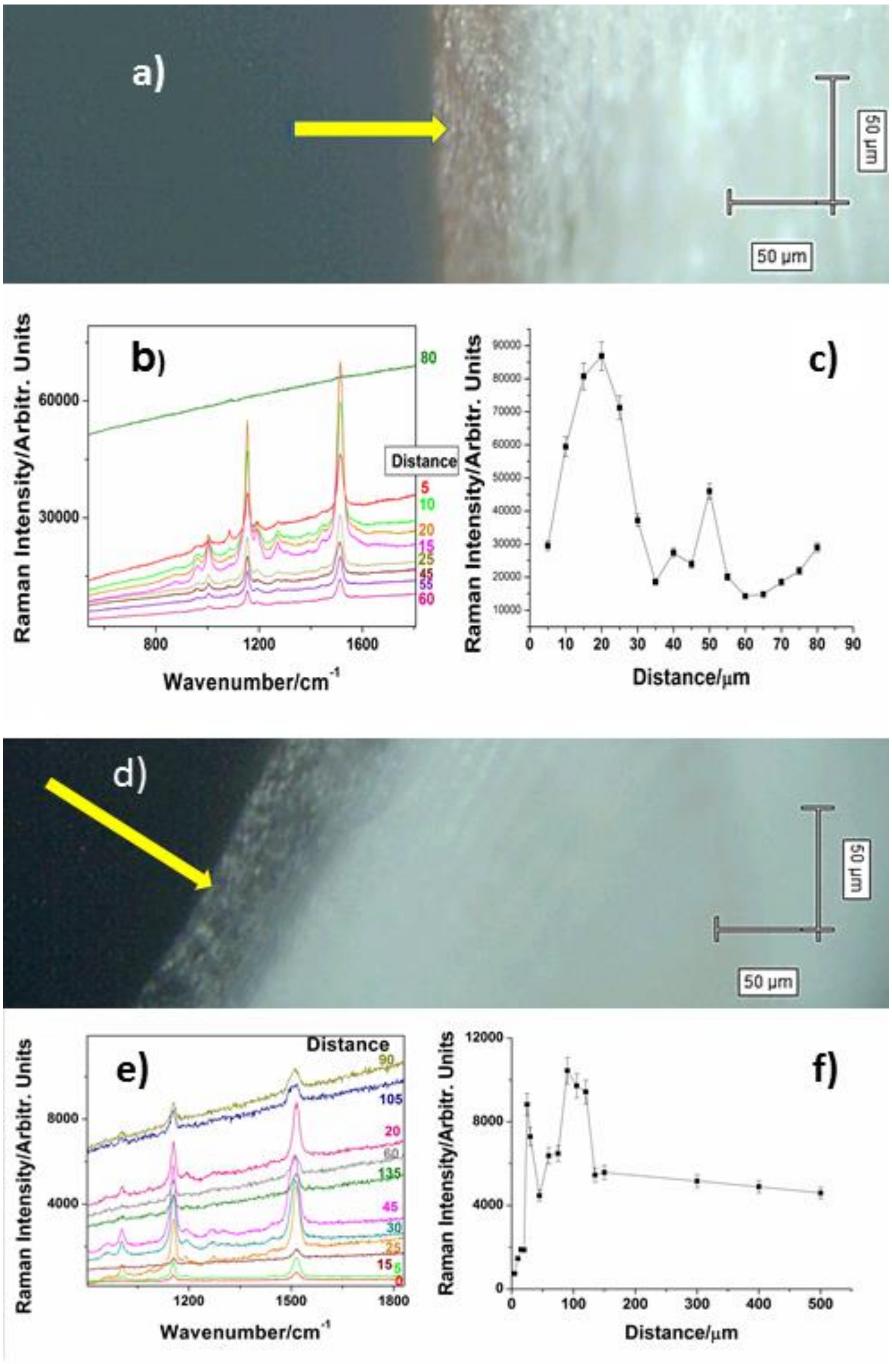


**Supplementary Fig. 7** Depth profiles of the ATX $v_1$ band intensity in the cross section of (a-c) *Callinectes sapidus* red claw shell, and (d-e) *Carcinus aestuarii* green claw shell. Micrographs (a, d) taken via Raman microscope during measurements show the cross-section planes of respective cuticles, featuring pale endocuticle, below the intensively colored exocuticle. Raman spectra (b, e) were taken along the transect line normal to surface, toward interior, as long as any notable signal-to noise was observed, using the 532 nm excitation line. Carotenoid content, measured by intensity of the $v_1$ Raman band (c, f), displays a variable distribution with a maximum at 10 to 20 µm below the cuticle surface in both shell types, followed by another peak at the depth of about 50 µm in red and 100 µm in green shell. The data suggest lower, but consistent carotenoid presence towards endocuticle inner layer. Error bars show the standard deviation.

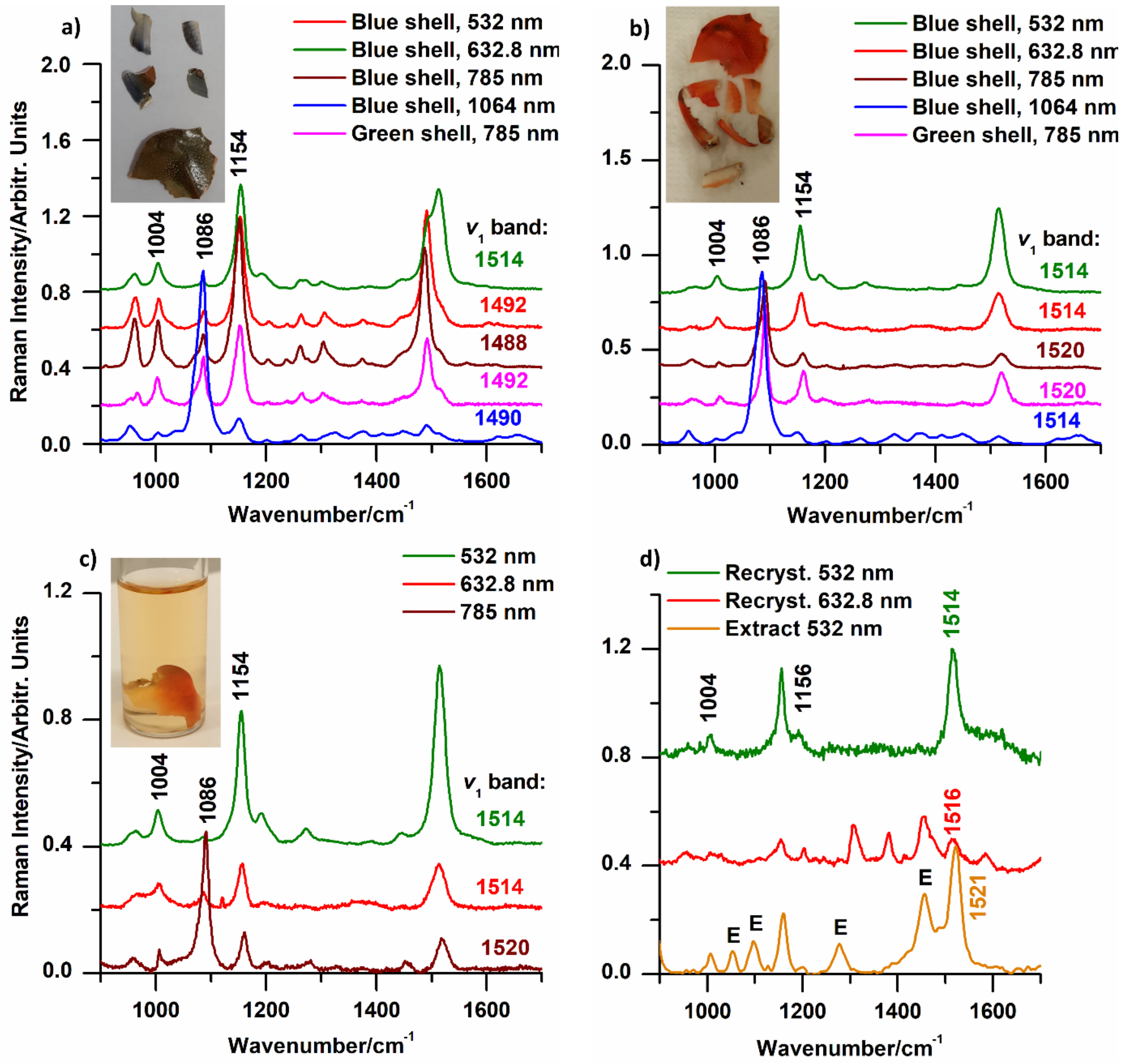


**Supplementary Fig. 8** Raman spectra (**b**ackground-subtracted, averaged) of *C. Sapidus* blue cuticle a) in native state and b) after boiling in hot water (100 °C) for 30 s. Note the disappearance of the ncb-ATX Raman mode at 1492 cm$^{-1}$ in Raman spectra collected from boiled relative to native shells, indicating complete denaturation of AXT-crustacyanin complexes followed by change of shell color from blue to orange (respective insets); c) native blue shell after 14 days of extraction in ethanol; note the absence of the mode assigned to ncb-ATX in crustacyanin and pink-yellowish color of astaxanthin-enriched ethanol extract in the inserted photo; d) shows that ethanol solvent was enriched with carotenoids, which could be again recrystallized after ethanol evaporation (E - ethanol bands).

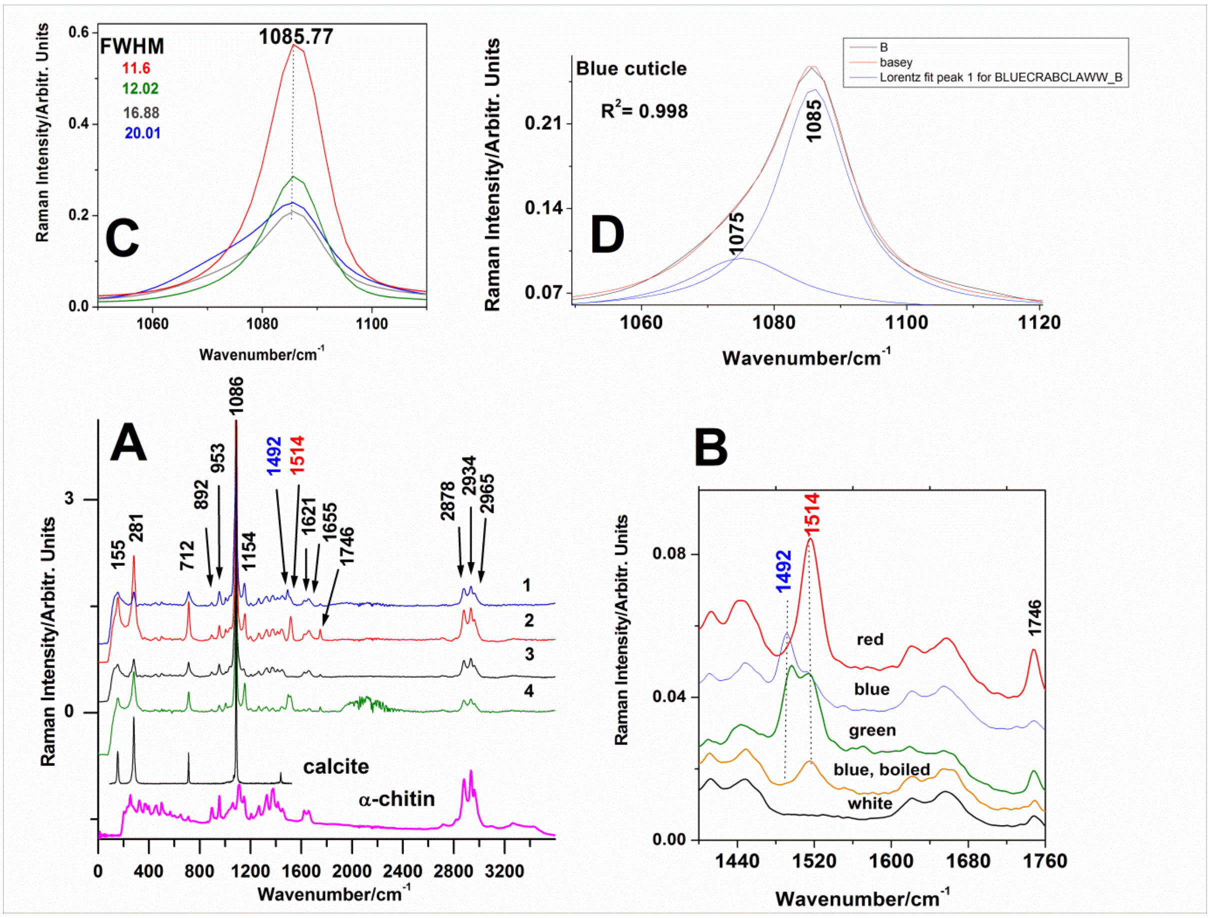


**Supplementary Fig. 9.** **A)** FT-Raman spectra collected from 1-blue, 2-red, 3-white and 4-green crab cuticle comparatively shown with the standard calcite and chitin spectra. B) Detail of the 1400-1780 $cm^{-1}$ range with the C=C mode in ATX at 1514 $cm^{-1}$ and ncb-ATX at 1492 $cm^{-1}$ in each coloured cuticle, along with the carbonate $2\nu_2$ mode at 1746 $cm^{-1}$. Note the disappearance of the 1492 $cm^{-1}$ band in the blue cuticle after immersion in boiling water. C) Details of the main carbonate mode (including its corresponding FWHM data for each cuticle color code); D) Lorentz fit of the broaden carbonate mode in blue shell, showing the crystalline calcite at 1085 and amorphous phase at 1075 $cm^{-1}$. Their area ratio was r= $area_{(1085)}/area_{(1075)}$=2.82. Excitation: 1064 nm, 350 mW.

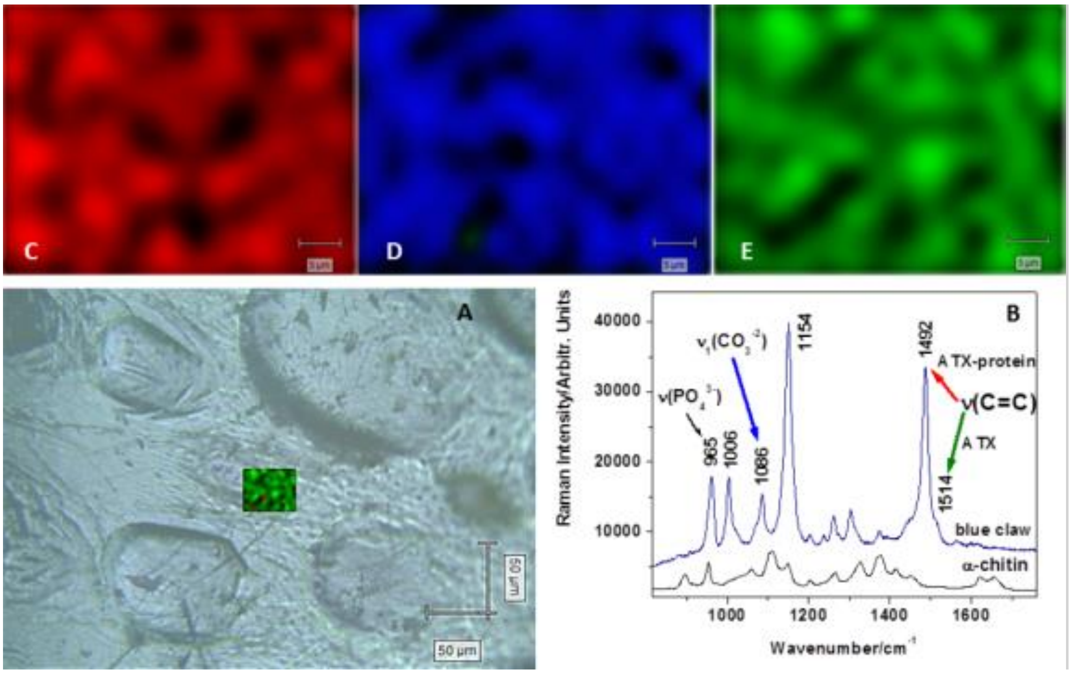


**Supplementary Fig. 10.** Mapping the signal-to-baseline distribution in blue cuticle (A) over an area of 50 μm x 40 μm exploiting its NIR-Raman spectrum excited with 785 nm (B) for carotenoproteins (C), calcite (D) and free ATX (E). Note the different pattern distribution of the respective bands intensity marked with red, blue or green arrays in B), suggesting high inhomogeneity of the chemical composition at micrometer scale and the interplay of the mineral and organic phase, well correlated with the morphology. Scale bar: 5 μm (C, D, E).

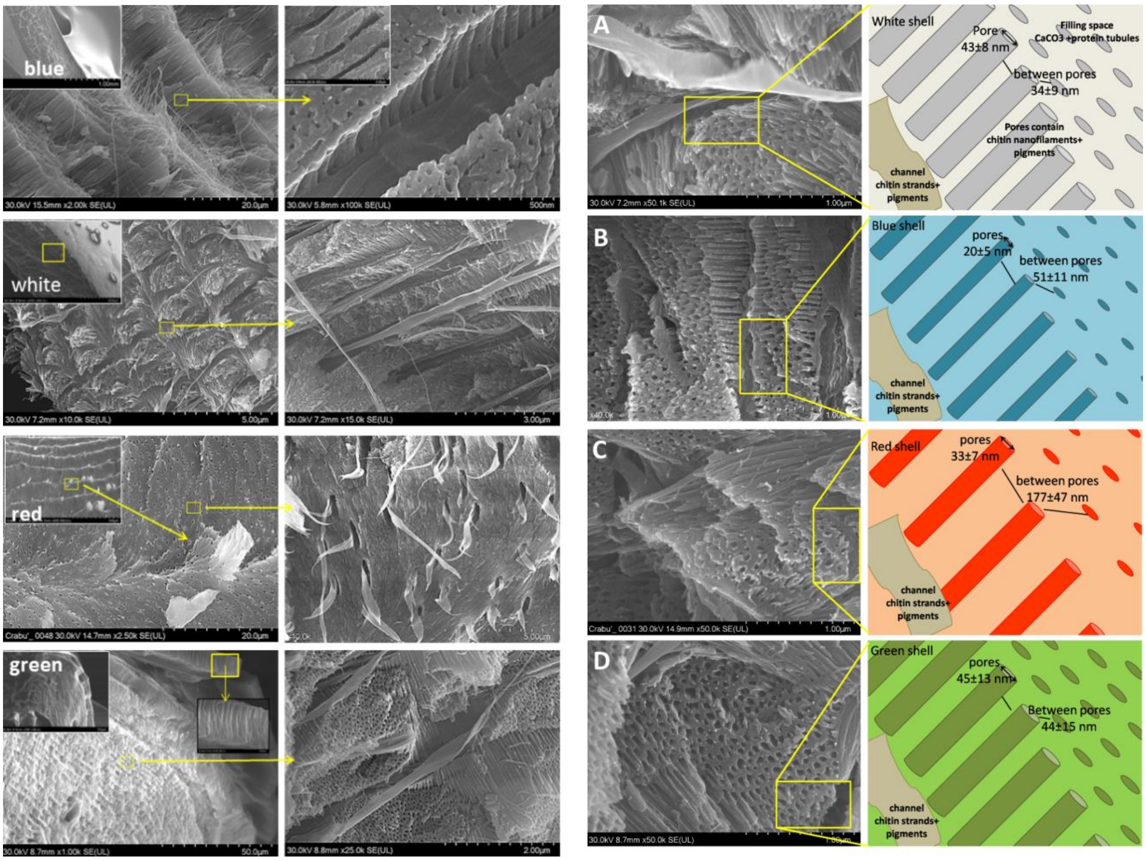


**Supplementary Fig. 11. Left:** Representative SEM images showing the morphology of the blue, white, red cuticle of *Callinectes sapidus* and green *Carcinus aestuarii* cuticle, as indicated. The yellow squares locate the higher magnification details highlighting the chitin-protein bundles, their corresponding canals and the regular arrays of nanopillars separated by pores in the ultrastructured canal walls, particularly illustrated in blue, white and green cuticle. Top view of the canals and the helical fibrils arising from them is highlighted in red cuticle. **Right:** Additional morphological details of crab cuticle nanoarchitecture from *C. sapidus* white (A), blue (B) and red (C), and *Carcinus aestuarii* green (D) claw shells. SEM images showing grating arrangement of pores and canals are shown along with their schematic representation, to highlight the pores diameter and inter-distances as super-grating, specific for each colour.

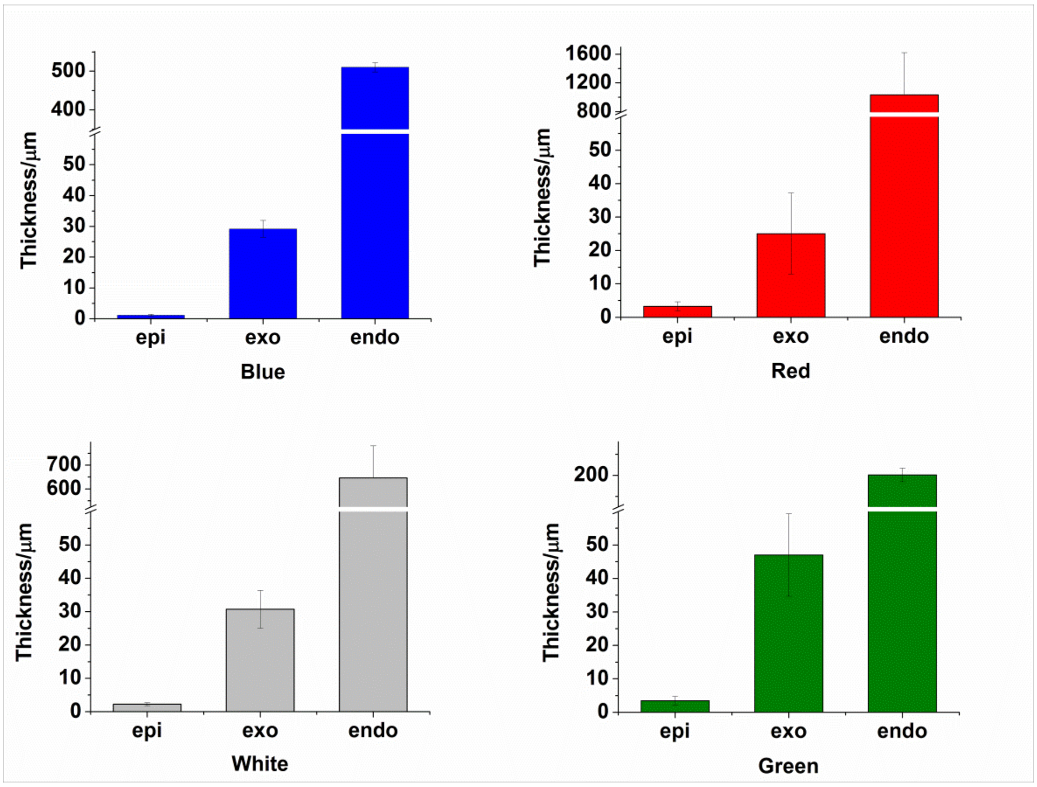


**Supplementary Fig.12** Comparative display of the averaged thickness of the cuticle layers, showing the thinnest epicuticle in blue shell. To calculate averaged data, at least 30 measurements have been conducted in distinct SEM images of layers. Error bars show the standard deviation.

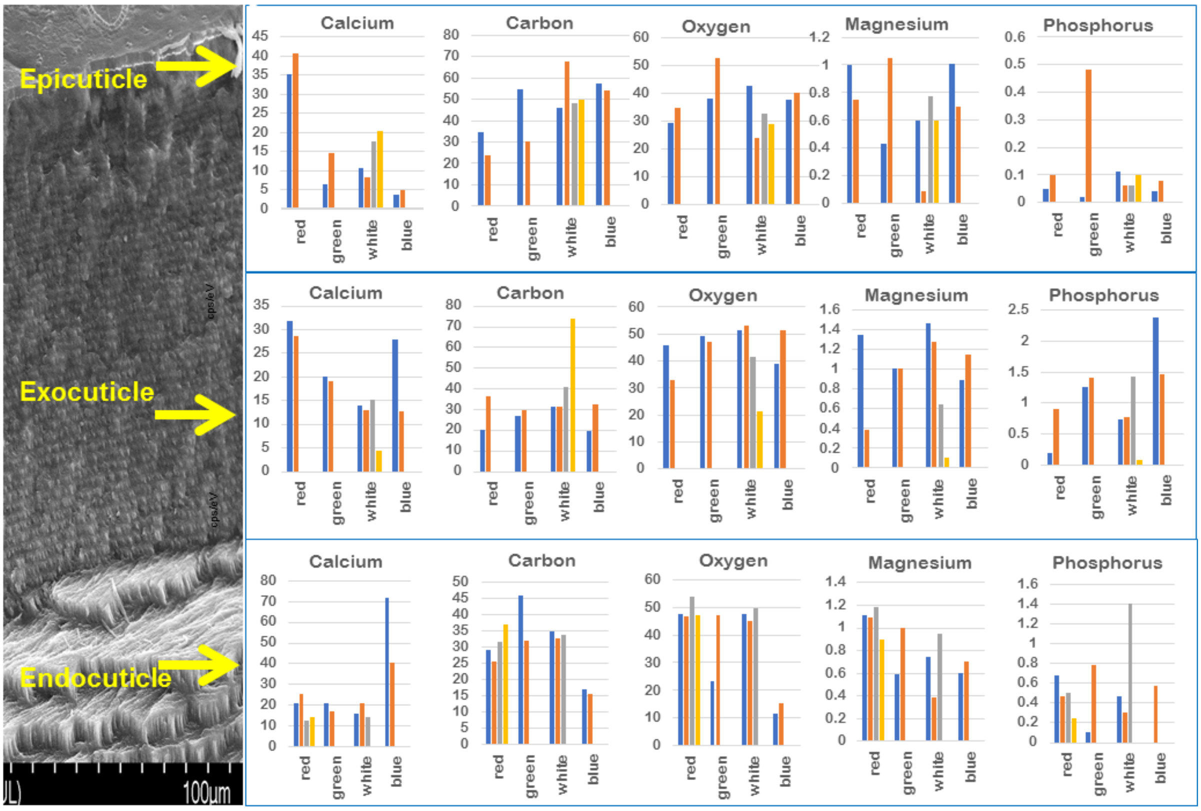


**Supplementary Fig. 13.** Relative mass fractions (wt%) of the five elements (Ca, C, O, Mg and P, as indicated) in crabs claw shells. EDX data were obtained from 2 to 4 scans corresponding to individual column color in epicuticle, exocuticle and endocuticle of *Callinectes sapidus blue, red and white* and *Carcinus aestuarii* green cuticle color, respectively.

**Supplementary Table S1.** Relative weight contribution (wt%) of all elements recorded in blue, red and white *C. sapidus* and green *C. aestuarii* claw cuticles extracted from the EDX measurements.

| **Cuticle color, layer** | **Green** | | | **Red** | | | **Blue** | | | **White** | | |
|---|---|---|---|---|---|---|---|---|---|---|---|---|
| **Element** | **endo** | **exo** | **epi** | **endo** | **exo** | **epi** | **endo** | **exo** | **epi** | **endo** | **exo** | **epi** |
| **C** | 38.705 | 28.26 | 42.45 | 30.555 | 28.38 | 28.965 | 15.64 | 26.005 | 55.75 | 33.646 | 44.29 | 52.97 |
| **O** | 40.04 | 48.315 | 45.21 | 48.885 | 39.26 | 31.96 | 26.89 | 45.135 | 38.78 | 47.443 | 41.915 | 31.947 |
| **Na** | 0.52 | 0.815 | 0.585 | 0.577 | 0.35 | 0.21 | 0.22 | 0.315 | 0.205 | 0.37 | 0.405 | 0.217 |
| **Mg** | 0.795 | 1.01 | 0.74 | 1.07 | 0.86 | 0.875 | 0.665 | 1.02 | 0.855 | 0.693 | 0.87 | 0.512 |
| **P** | 0.45 | 1.33 | 0.25 | 0.485 | 0.54 | 0.075 | 0.285 | 1.92 | 0.06 | 0.74 | 0.7475 | 0.082 |
| **K** | 0.2 | 0.21 | 0.065 | 0.062 | 0.17 | 0.065 | 0.27 | 0.24 | 0.035 | 0.14 | 0.0866 | 0.06 |
| **Ca** | 18.955 | 19.605 | 10.6 | 18.342 | 30.29 | 37.86 | 56.03 | 25.365 | 4.32 | 17.06 | 11.667 | 8.18 |
| **S** | | | | | | | | | | | 0.16 | |
| **Cl** | 0.34 | 0.46 | 0.1 | 0.07 | 0.29 | | | | | | | |

*Abbreviations*: endo – endocuticle, exo – exocuticle, epi - epicuticle.
